# The core MICOS complex subunit Mic60 has been substituted by two cryptic mitofilin-containing proteins in Euglenozoa

**DOI:** 10.1101/2025.01.31.635831

**Authors:** Shaghayegh Sheikh, Barbora Turpin Knotková, Corinna Benz, Marek Eliáš, Tomáš Bílý, Alexey Bondar, Martina Tesařová, Eva Stříbrná, Jiří Heller, Michael Meinecke, Hassan Hashimi

**Affiliations:** Institute of Parasitology, Biology Centre, Czech Academy of Sciences, České Budějovice, Czech Republic; University of South Bohemia, Faculty of Science, České Budějovice, Czech Republic; Heidelberg University Biochemistry Center (BZH), Heidelberg, Germany; Department of Biology and Ecology, Faculty of Science, University of Ostrava, Ostrava, Czech Republic; Institute of Entomology, Biology Centre, Czech Academy of Sciences, České Budějovice, Czech Republic

**Author notes:** Department of Medical Biochemistry and Biophysics, Division of Molecular Metabolism, Karolinska Institutet, Biomedicum, 171 65, Solna, Sweden (Shaghayegh Sheikh).

## Abstract

Cristae enclose respiratory chain complexes, making them the bioenergetic subcompartments of mitochondria. The MICOS complex is among the inducers of membrane curvature needed for crista formation. Resembling the respiratory chain complexes, MICOS is organized around a core protein, the mitofilin-domain bearing Mic60, that was inherited from the alphaproteobacterial progenitor of mitochondria. Extant alphaproteobacteria express Mic60 to form their own bioenergetic subcompartments, demonstrating the permeance of Mic60’s form and function during prokaryotic and eukaryotic evolution. Yet, unlike virtually all aerobic eukaryotes, Mic60 is not encoded within the genomes of the multifarious protists that comprise the phylum Euglenozoa, including trypanosomes. Here, we show that Mic60 has been replaced in euglenozoans by two cryptic mitofilin domain-containing MICOS subunits, Mic34 and Mic40. Contrasting alphaproteobacterial and mitochondrial Mic60, these are not integral membrane proteins. Mic34 and Mic40 are as diverged from each other as both are to canonical Mic60. Reverse genetics revealed they are intertwined with the oxidative protein folding pathway required for mitochondrial–and crista–biogenesis, veiling a potential membrane remodeling role. Nevertheless, Mic34 binds phospholipid bilayers *in vitro*. Mic34 and Mic40 heterologous expression remodels gammaproteobacterial cytoplasmic membranes, like Mic60. Unexpectedly, Mic34 overexpression elaborates the simplified tubular mitochondrion of a *Trypanosoma brucei* life cycle stage with repressed oxidative phosphorylation. Furthermore, this activity was ablated by mutations to Mic34’s mitofilin domain that correspond to essential motifs found in yeast Mic60’s mitofilin domain. Thus, the mitofilin protein family is more diverse than originally supposed, with two of its structurally most divergent members altering the core of euglenozoan MICOS.

## INTRODUCTION

Cristae are the bioenergetic subcompartments of mitochondria that house the molecular machinery for cellular respiration (Pánek et al. 2020). Their biogenesis involves the extreme bending of the mitochondrial inner membrane (IM) by dedicated–and usually multiprotein– molecular machineries (Barbot and Meinecke 2016; Daumke and van der Laan 2025). Rows of ATP synthase dimers are responsible for positive curvature at cristae rims (Kühlbrandt 2019) whereas the Mitochondrial contact site and Cristae Organizing System (MICOS) complex forms both crista junctions, constricted zones of negative curvature (Colina-Tenorio et al. 2020). Additionally, MICOS forms contact sites between the IM and outer membrane.

While ATP synthase dimerization appears to be a eukaryotic invention (Kühlbrandt 2019), the keystone MICOS subunit Mic60 is of more ancient origin, being directly inherited from the alphaproteobacterial symbiont that gave rise to mitochondria (Muñoz-Gómez et al. 2015; Huynen et al. 2016). Mic60’s function in the formation of bioenergetic subcompartments and double membrane contact sites is conserved in extant alphaproteobacteria (Muñoz-Gómez et al. 2023), supporting the hypothesis that the cenancestor of mitochondria and alphaproteobacteria already possessed the requisite machinery to form cristae (Muñoz-Gomez et al. 2017). Indeed, the shared properties of eukaryotic and prokaryotic Mic60 most parsimoniously imply that their common ancestor had the capacity to remodel their inner membranes into crista-like subcompartments (Tarasenko et al. 2017). Also, like the respiratory chain complexes that are embedded in cristae, MICOS is also assembled around a proteinaceous core inherited from the alphaproteobacterial symbiont (Roger et al. 2017). Core architectures are necessarily more conserved than the surrounding shell of often lineage-specific subunits to stabilize the functionality of these protein complexes (Prokopchuk et al. 2023).

The canonical domain architecture of Mic60 (Muñoz-Gómez et al. 2015; Huynen et al. 2016) forms a tertiary structure that allows the protein’s homotetramerization, which has been proposed to tubulate membranes (Bock-Bierbaum et al. 2022). A transmembrane domain (TMD) embeds its N-proximal segment into the IM. A central coiled coil (CC) domain provides an interaction interface bringing together two antiparallelly oriented Mic60 homodimers into a tetramer. The C-terminal mitofilin domain, made up of eight alpha helices, directly interacts with phospholipids on the opposite side of a tubulated membrane. Via this mechanism, Mic60 has been proposed to be the primary factor in crista junction constriction (Stephan et al. 2020).

Given its central role in crista biogenesis, Mic60 is ubiquitously encoded in the genomes of aerobic eukaryotes, with only a few exceptions (Muñoz-Gómez et al. 2015; Huynen et al. 2016). Flagellated protists belonging to the phylum Euglenozoa represent one such anomaly. Euglenozoans are a sub-clade within the eukaryotic supergroup Discoba (Burki et al. 2020). This phylum is named after Euglenida, a subdivision containing free-living, mostly freshwater protists (Fig. 1A). Two sister clades branch next to Euglenida: Diplonemea, a group of predominantly marine flagellates, and Kinetoplastea (or Kinetoplastida), named after the kinetoplast, a massive network of mitochondrial DNA that comprises the organelle’s single genome (Kostygov et al. 2021). A species-poor group Symbiontida forms a fourth euglenozoan lineage (Lax et al. 2021), but it is not considered further in this study due to limited sequence resources.

**Fig. 1.**
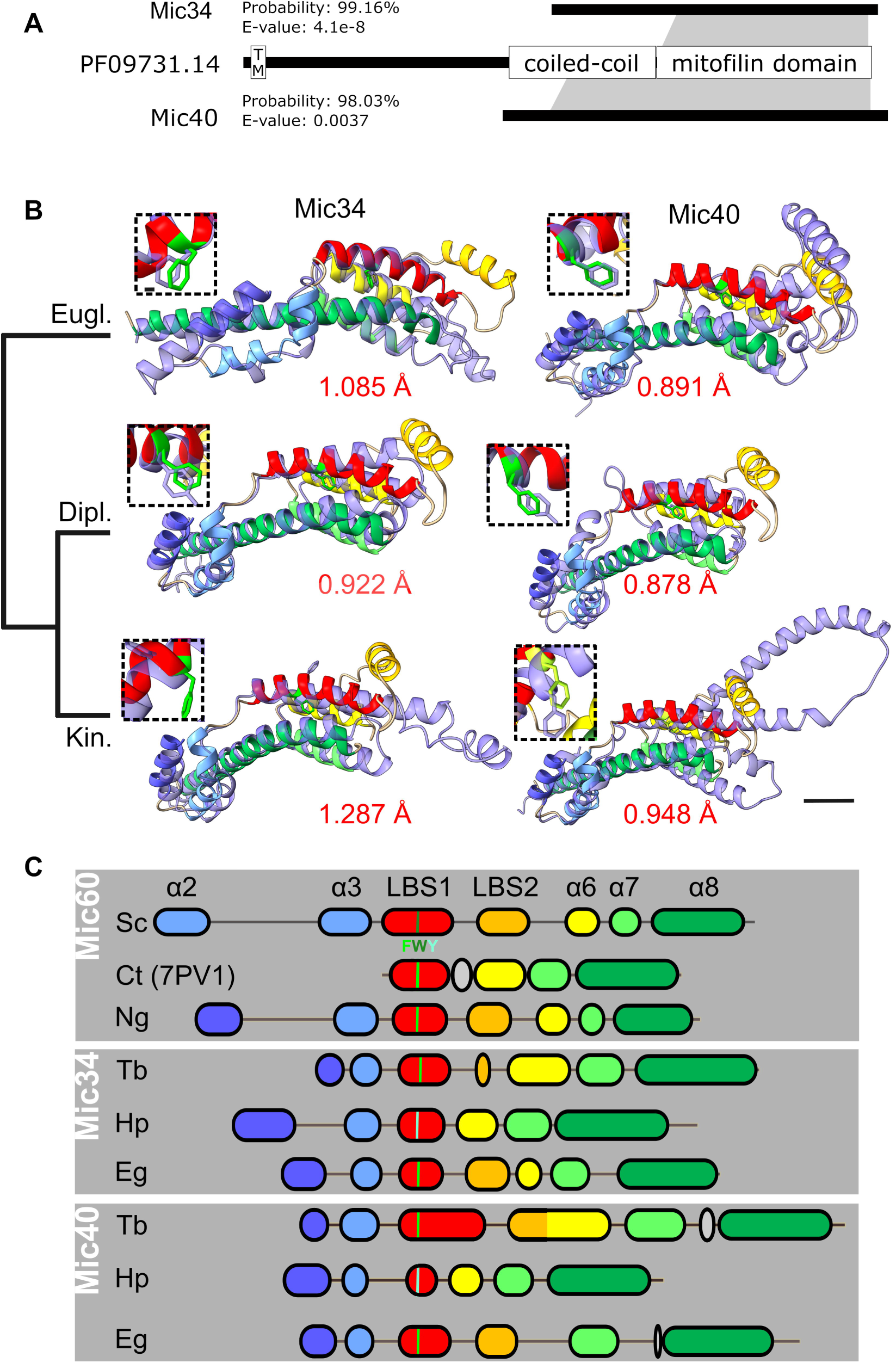
The N-termini of Mic34 and Mic40 retain a mitofilin domain. **(A)** Schematic representation of HHpred results using representative profile HMMs. The searches detected statistically significant matches (the respective values indicated in the figure) between both Mic34 and Mic40 and the Pfam family PF09731.14 (*i.e.*, Mic60), with the specific matching regions delimited by the grey zones. The three proteins are drawn to scale according to the reference proteins for each HMM (arbitrarily Mic34 and Mic40 from *T. brucei* and *Dictyostelium discoideum* Mic60). For Mic60 (PF09731.14) three different structural/functional regions are delimited, including a TMD (TM), a coiled-coil region (boundaries as reported (Hessenberger et al. 2017) for *C. thermophila* Mic60), and the mitofilin domain as recently redefined (Benning et al. 2025). **(B)** The AlphaFold2 predicted structures of the putative mitofilin domains of Mic34 and Mic40 (light purple) aligned to the predicted mitofilin domain of discoban *N. gruberi* Mic60 (alpha helices colored as in B). The root-mean-square distance (RMSD) of the alignments is given in red below the model. Scale bar, 10 Å. Dashed-boxed insets show alignment of conserved LBS1 aromatic amino acid. Scale bar, 1 Å. Major euglenozoan clades depicted as in (Kostygov et al. 2021): Eugl., Euglenida; Dipl., Diplonemea; Kin., Kinetoplastea. **(C)** Map of the mitofilin alpha-helical motifs from yeast and *N. gruberi* (Ng) Mic60, plus representative Mic34 and Mic40 from the three clades in A. Mitofilin alpha helices are color coded and correspond to names on top. The conserved aromatic amino acid residue of LBS1 is color coded based on the key below the uppermost LBS1 (F, phenylalanine; W, tryptophan; Y, tyrosine). Abbreviations: α, alpha helix; Ct, *C. thermophilum*; Eg, *E. gracilis*; Hp, *H. phaeocysticola*; LBS, lipid binding site; Sc, *S cerevisiae*; Tb, *T. brucei*. The PDB accession number for the solved CtMic60 mitofilin domain structure (Bock-Bierbaum et al. 2022), in which LBS2 was necessarily deleted, is given in parentheses.

Until it was isolated from the kinetoplastid *Trypanosoma brucei* (Kaurov et al. 2018), the MICOS complex was only known from animals and yeast, grouped together in the diverse Opisthokonta clade (Burki et al. 2020). *T. brucei* MICOS has some commonalities with opisthokont MICOS, such as having the other central subunit, Mic10. While *T. brucei* MICOS differs in having two Mic10 paralogs, one paralog associates with ATP synthase dimers (Cadena et al. 2021), like in yeast (Rampelt et al. 2022). But *T. brucei* MICOS also has traits not observed in opisthokont MICOS. One example is that half of the subunits assemble into a soluble subcomplex within the intermembrane space (IMS; Fig. 2C) (Kaurov et al. 2018; Eichenberger et al. 2019). Furthermore, it seems to be directly involved in a still unknown aspect of the Mitochondrial Intermembrane Space Assembly (MIA) pathway (Kaurov et al. 2018; Kaurov et al. 2022), which facilitates the oxidative folding of proteins imported into the IMS (Mordas and Tokatlidis 2015). Here, we address another trait that sets *T. brucei* MICOS apart from MICOS in virtually all other eukaryotes. As predicted by the aforementioned bioinformatic screen, the isolated *T. brucei* MICOS lacked a canonical Mic60. At the time of discovery, a subunit bearing a superficially similar domain architecture, but lacking the characteristic mitofilin domain, was annotated as a putative Mic60 (Kaurov et al. 2018).

**Fig. 2.**
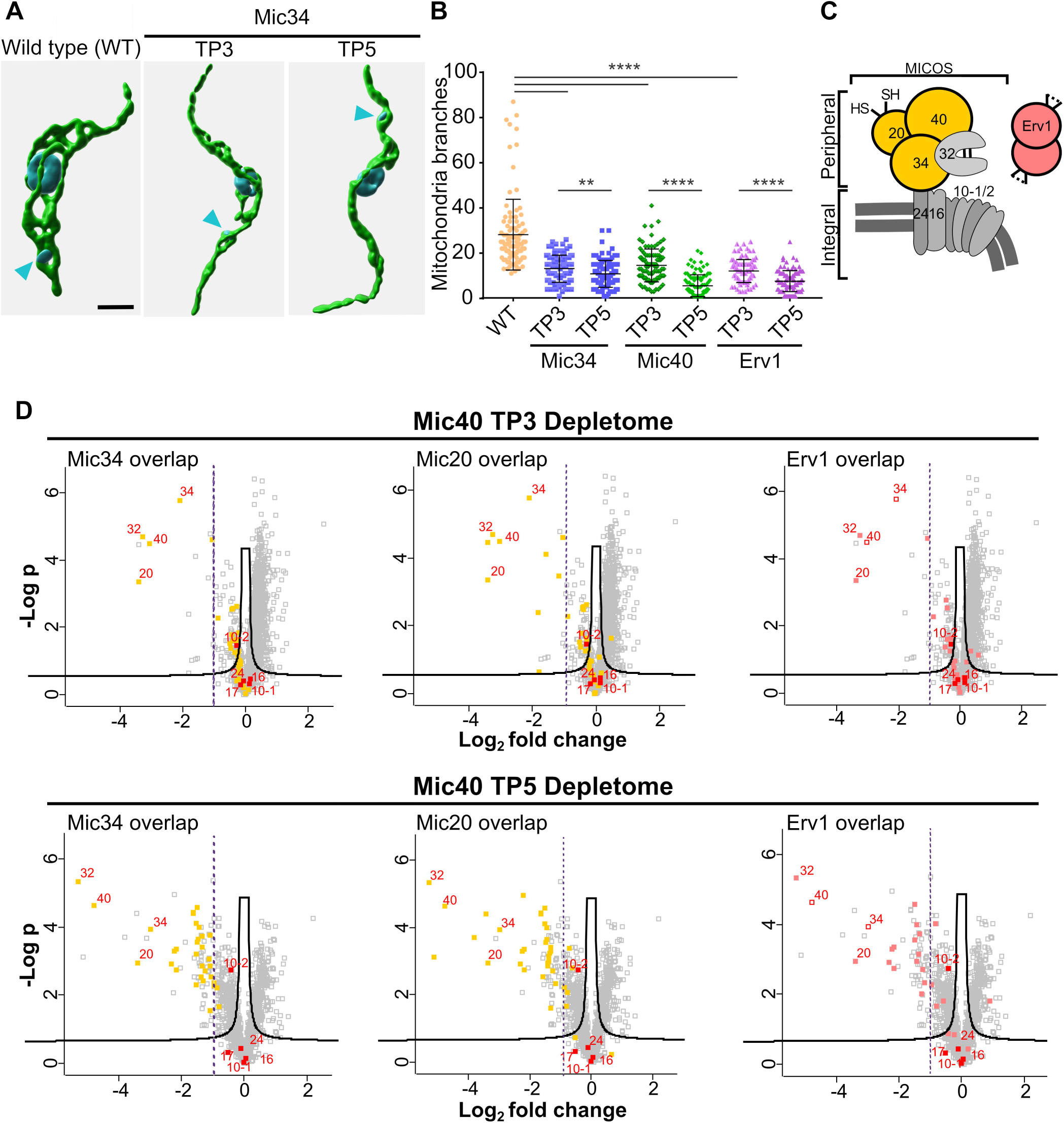
Mitochondrial remodeling by Mic34 and Mic40 seems to be coupled with their involvement in the mitochondrial intermembrane space assembly pathway. **(A)** Representative 3D reconstruction of mitochondria in wild type (WT) and Mic34 knockdown *T. brucei* cells at time points (TPs) 3 and 5 days after Dox induction. Mitochondria labeled by an anti-HSP70 antibody are in green; the nucleus and kinetoplast DNA (arrowhead) are in blue. Scale bar, 1 µm. **(B)** Scatter plot showing the number of mitochondrial branches per cell (n=100) in WT plus Mic34, Mic40, and ERV1 RNAi at TP 3 and 5 of doxycycline induction. Data are presented as individual values with their mean and standard deviation indicated by bar and whiskers. Statistical significance: **p = 0.01 and ****p < 0.0001. **(C)** Schematic representation of the MICOS complex and ERV1. Peripheral MICOS subunits whose depletomes were determined are colored in yellow. Integral membrane components are shown in gray. The lipid bilayer is represented as a double gray line. The ERV1 dimer is depicted in red. **(D)** Volcano plots of Mic40 depletomes measured at TP3 (top row) and TP5 (bottom) of RNAi induction. Each row shows the same Mic40 TP3 and TP5 depletome data colored to highlight overlap with the depletomes of (left to right) Mic34 (Eichenberger et al. 2019), Mic20 (Kaurov et al. 2018; Kaurov et al. 2022), and Erv1 (Peikert et al. 2017). Points corresponding to MICOS subunits are individually labelled as in C. X-axis, Log_2_-transformed fold change of Mic40 RNAi over WT; y-axis, -log of P-value of measurement from triplicates.

However, here we present *in silico* and experimental evidence that two other MICOS subunits named Mic34 (Uniprot ID: Q57ZZ1) and Mic40 (Q57YH2) truly represent cryptic mitofilin family proteins. Thus, we henceforth will refer to the originally misannotated Mic60 (Q38DZ0) as Mic24. This discovery has expanded the mitofilin proteins to include members that implement the eponymous domain unconventionally.

## RESULTS

### The soluble MICOS subunits Mic34 and Mic40 are paralogs that each contain a mitofilin domain

A *bona fide* ortholog of Mic60 was not identified in the MICOS complex isolated from *T. brucei* (Kaurov et al. 2018). However, when the largest *T. brucei* Mic34 and Mic40 MICOS subunits were searched against the Pfam-A (v.37) database using HHpred (Söding et al. 2005), which implements sensitive protein homology detection based on pairwise comparison of profile hidden Markov models (HMMs), both retrieved the family PF09731.14, *i.e.* mitofilin as the best hit with convincing statistical significance: 98.08% probability and E-value of 0.0056 for Mic34, and 96.77% probability and E-value of 2.2 for Mic40. We then built a more representative profile HMM based on Mic34 and Mic40 multiple sequence alignments (MSAs) of a phylogenetically comprehensive set of homologs retrieved from all major Euglenozoa groups, i.e. Kinetoplastea, Diplonemea, and Euglenida (supplementary dataset S1A, Supplementary Material online). The improved taxonomic coverage of the query MSA increased the sensitivity of the homology detection by HHpred and reinforced the credibility of the matches to mitofilin for Mic34 and Mic40 by increasing the probabilities to 99.16% and 98.03%, respectively, and lowering the E-values to 2.3e-7 and 0.021, respectively (supplementary figs. S1 and S2, Supplementary Material online). In contrast, a HHpred search with *T. brucei* Mic24 (formerly ‘Mic60’, see above) did not retrieve Mic60 or the mitofilin domain even among all 250 best hits displayed by default. This did not change when we used as queries custom alignments of a broader set of kinetoplastid or euglenozoan Mic24 homologs (listed in supplementary dataset S1B, Supplementary Material online). Comparison of profile HMMs derived from MSAs of the kinetoplastid and euglenid Mic24 sequences revealed a region of homology that was restricted to a previously delimited C-terminal α-helical region (Kaurov et al. 2018). We conclude that this represents a novel Mic24-specific domain without homologs discernible by HHpred. Notably, in contrast to Mic34 and Mic40 with ubiquitous occurrence in euglenozoans, no Mic24 homologs were identified in diplonemids and the endosymbiotic kinetoplastids of the genus *Perkinsela*.

Inspection of the HHpred Mic34 and Mic40 alignments to the mitofilin protein family entry showed that the Mic60 region detected to be homologous to one or the other euglenozoan protein differs (Fig. 1A). For the Mic34-Mic60 comparison the match was restricted to a C-terminal part of the former and a region of the latter exactly corresponding to the ‘mitofilin domain’ as delineated by Benning et al. (2025). The match between Mic40 and Mic60 extended further upstream to include a larger part of the Mic60’s central coiled-coil (CC) region. This alignment covered almost the full Mic40 protein, except for a short N-terminal region, which corresponds to a part of Mic40 not detected by mass spectroscopy in a previous study, implying it potentially is not part of the mature protein (Kaurov et al. 2018). To further illuminate the relationship between Mic60 and euglenozoan Mic34 and Mic40, we focused on the mitofilin domain apparently shared by all three proteins. For this purpose, we collected a reference set of Mic60 sequences enriched with proteins from eukaryotes most closely related to Euglenozoa (i.e., other phyla of the supergroup Discoba; see supplementary dataset S1C, Supplementary Material online) and created a custom alignment of their mitofilin domain. Pairwise profile HMM-HMM comparisons of the different mitofilin domains using HHpred indicated a higher sequence similarity in the case of Mic34-Mic60 comparison than for the Mic40-Mic60 comparison (the respective alignment scores being 93.09 and 72.52, respectively). Notably, an analogous comparison of the Mic34 and Mic40 mitofilin domain HMMs yielded an alignment restricted to their C-terminal parts (and hence with a much lower score, 47.38 or 49.55 depending on which domain is used as a query). Phylogenetic analysis of the mitofilin domain sequences divided them into the three expected groups, albeit with low bootstrap support for each clade, and confirmed the mutually high divergence of Mic34 and Mic40 sequences (manifested by their branch length; supplementary fig. S3, Supplementary Material online). The most parsimonious scenario explaining the origin of Mic34 and Mic40 starts with a canonical *mic60* gene in the euglenozoan ancestor followed by its duplication into the eventual Mic34 and Mic40 paralogs. Subsequent rapid evolution of the two paralogs diverged them considerably from each other, both in the N-terminal region of the encoded protein as well as in the mitofilin domain. Owing to the presence of both Mic34 and Mic40 orthologs in all three major euglenozoan lineages, all these events must have happened before their divergence.

The mitofilin domain is the most conserved primary structure of prokaryotic and eukaryotic Mic60 proteins (Muñoz-Gómez et al. 2015; Huynen et al. 2016). To bolster confidence that Mic34 and Mic40 are cryptic mitofilin proteins, we employed AlphaFold2 predictions of the mitofilin domains starting from its second alpha helix (α2) (Benning et al. 2025) found in the three representatives from each euglenozoan group, namely *T. brucei* (Kinetoplastea), *Hemistasia phaeocysticola* (Diplonemea), and *Euglena gracilis* (Euglenida) (Fig. 1B). These tertiary structures were aligned to the mitofilin domain of discoban *Naegleria gruberi* Mic60, which overlaps the theoretical *Saccharomyces cerevisiae* and the experimentally solved *Thermochaetoides thermophila* (hereafter referred to by its previous name *Chaetomium thermophilum*) (Bock-Bierbaum et al. 2022) mitofilin domains with a root-mean-square distance (RMSD) of <1 Å; *N. gruberi*’s mitofilin aligned to each fungal one equivalently as the two fungal structures did to each other (supplementary fig. S4A, Supplementary Material online). Expectedly, the Mic34 mitofilin domains of the three euglenozoan representatives are well aligned (supplementary fig. S4B, Supplementary Material online).

A linear map of Mic34 and Mic40’s mitofilin domain alpha helices α2-α8 (Benning et al. 2025) was derived from the high confidence predicted structures (Fig. 1C) and multiple sequence alignments used to make the aforementioned profile HMM (Fig. 1A) and phylogenetic tree (supplementary fig. S3, Supplementary Material online). The extended loop between α2 and α3 in Mic60 *S. cerevisiae* (and to a lesser extent *N. gruberi*) is considerably reduced in Mic34 and Mic40. Lipid binding site (LBS) 1, which is necessary for optimal lipid binding and yeast cristae junction formation (Hessenberger et al. 2017), was present in all examined Mic34 and Mic40. In almost all structural alignments, a highly conserved and functionally important aromatic amino acid (Muñoz-Gómez et al. 2023) within LBS1 perfectly overlapped (Fig. 1B; supplementary fig. S4A<u>, Supplementary Material online</u>). This phenylalanine is staggered in *T. brucei* compared to all the other LBS1 structures examined (Fig. 1B; supplementary fig. S4B, Supplementary Material online) because it is shifted by one amino acid in the kinetoplastid primary structures (supplementary fig. S7F<u>, Supplementary Material online</u>). Discoban LBS1 appears to be an amphipathic helix with the hydrophobic face oriented like fungal LBS1; a notable exception is *T. brucei* Mic34, whose hydrophobic face is shifted ∼90° counterclockwise (supplementary fig. S4D<u>, Supplementary Material online</u>).

The adjacent LBS2 (α5) was much more variable, consistently being the least reliably predicted motif that aligned poorly to the reference *N. gruberi* mitofilin domain (supplementary fig. S4C<u>, Supplementary Material online</u>). Furthermore, it is missing in *H. phaeocysticola* Mic34 and Mic40. The orientation of the last three alpha helices α6-8 is well conserved in all three representative Mic34 proteins. In Mic40, this is a more malleable feature, with *T. brucei* and *E. gracilis* having a shorter alpha helix downstream of α8, and α6 fused to LBS2 or missing, respectively (Fig. 1C). In summary, the mitofilin domain structure of Mic34 appears to be more similar to canonical Mic60 than Mic40.

### Mic34 and Mic40 are involved in the mitochondrial intermembrane space assembly pathway that affects mitochondrial morphology

To verify *in silico* predictions that Mic34 and Mic40 are cryptic mitofilins, we examined how their depletion in procyclic stage *T. brucei* (PCF), which bears mitochondria with mature cristae that house an active electron chain (Bílý et al. 2021), affects the organelle’s morphology. These inducible-RNAi cell lines were previously characterized (Kaurov et al. 2018; Eichenberger et al. 2019). Here, we recapitulated the expected growth inhibition upon RNAi silencing of Mic34 and Mic40 (supplementary fig. S5A<u>, Supplementary Material online</u>). Qualitative (Fig. 2A; supplementary fig. S5B<u>, Supplementary Material online</u>) and quantitative (Fig. 2B) assessment of mitochondrial complexity showed that branching was reduced at 3- and 5-day time points (TP3 and TP5) after RNAi induction, with the organelle exhibiting a tubular appearance in the latter. *S. cerevisiae mic60* null mutants also exhibit alternations to the mitochondrial network (Rabl et al. 2009).

Interpretation of the observed morphological defects may be confounded by downregulation of proteins imported into the IMS by the MIA pathway, which has been observed upon Mic34 silencing (Eichenberger et al. 2019). This role corresponds with the IMS localization of Mic34 and Mic40 (Kaurov et al. 2018).To see if Mic40 depletion has the same effect on IMS proteins as Mic34, we first verified that the MIA substrate Tim9 (Peikert et al. 2017) was downregulated during the time course of RNAi induction (supplementary fig. S5C, <u>Supplementary Material online</u>). To gain a comprehensive view on the cohort of proteins that may rely on Mic40 for their import, we performed a mitoproteome-wide analysis (*i.e.* depletome) of Mic40 TP3 and TP5 relative to wild type *T. brucei* (Fig. 2C-D). The MICOS subcomplex subunits residing completely in the IMS peripherally to the IM, were downregulated prior to reduction of MIA substrates (Fig. 2C-D). A similar phenotype was observed upon Mic34 silencing (Eichenberger et al. 2019). Indeed, the Mic40 depletome overlapped with those of Mic34 (Fig. 2D, left) and Mic20 (Fig. 2D, middle), a thioredoxin-like subunit of the peripheral, IMS-localized MICOS subcomplex (Kaurov et al. 2018; Kaurov et al. 2022). Thus, Mic40 also plays an integral part in IMS protein import by MIA.

Returning to the question of whether IMS import defects may underlie the reduction in mitochondrial complexity seen in Mic34 and Mic40 knockdown cells, we measured the degree of branching in TP3 and TP5 Erv1 RNAi PCF (Fig. 2B; supplementary fig. S5B, <u>Supplementary Material online</u>). Erv1 is a sulfhydryl oxidase that plays an essential role in the MIA pathway (Mordas and Tokatlidis 2015; Peikert et al. 2017), and is not a component of *T. brucei* MICOS (Kaurov et al. 2018; Turra et al. 2021). Erv1 depletion phenocopies those of Mic34 and Mic40 to the same degree Erv1 depletion’s effect on mitochondrial complexity was previously reported based on qualitative observations (Niemann et al. 2013; Haindrich et al. 2017). The Mic34 and Mic40 depletomes overlap with Erv1’s (Peikert et al. 2017). It should be noted that peripheral MICOS subunits were also downregulated upon Erv1 silencing (Peikert et al. 2017), possibly leading to the observed morphological alteration. Consistent with this hypothesis, these subunits were by far the most affected at TP3 (Fig. 2D), when simplification of the mitochondrion was first observed during Mic40 downregulation (Fig. 2A-B). However, while these results do not rule out that Mic34 or Mic40 are directly involved in membrane remodeling, we still cannot decouple this function from its established but still unexplained role in the MIA pathway.

### Mic34 binds to phospholipids *in vitro*

First, we decided to test if Mic34 and Mic40 can deform liposomes *in vitro*, as has been demonstrated for Mic60 (Hessenberger et al. 2017; Tarasenko et al. 2017). For this experiment, recombinant *T. brucei* Mic34 and Mic40 bearing N-terminal His-tags (rMic34 and rMic40, respectively) were expressed in *Escherichia coli*, where they sequestered into inclusion bodies (supplementary fig. S6A-B, Supplementary Material online). After their purification by sequential affinity and size exclusion chromatography, rMic34 and rMic40 were solubilized with varying concentrations of the mild detergent n-dodecyl β-d-maltoside (DDM). While the majority of rMic34 proved to be soluble in 0.1% DDM, rMic40 was poorly solubilized in any of the tested DDM concentrations (supplementary fig. S6C-D, Supplementary Material online). Thus, we excluded rMic40 from further analysis.

Next, 0.1% DDM solubilized rMic34 was loaded onto large unilamellar vesicles (LUVs) comprised of IM phospholipids (Barbot et al. 2015; Tarasenko et al. 2017), followed by DDM removal. A fraction of rMic34 was carbonate extracted from the LUVs (Fig. 3A), which is the expected behavior of the IMS-periphery localized Mic34 ((Kaurov et al. 2018) and also see below). Surprisingly, most rMic34 acted as an integral protein in this biochemical assay. To eliminate the possibility of DDM-mediated rMic34 insertion into the membrane, we performed a Proteinase K protection assay to see if Mic34 is binding the exterior of the LUV (Fig. 3B). The protease completely degraded rMic34 in the liposome sample even in the absence of the membrane-disrupting detergent sodium dodecyl sulfate (SDS). This mimicked the behavior of the *S. cerevisiae* rMic19, which serves as a negative control of phospholipid binding due to an absence of this *in vitro* activity (Hessenberger et al. 2017), and thus remains exposed to Proteinase K.

**Fig. 3.**
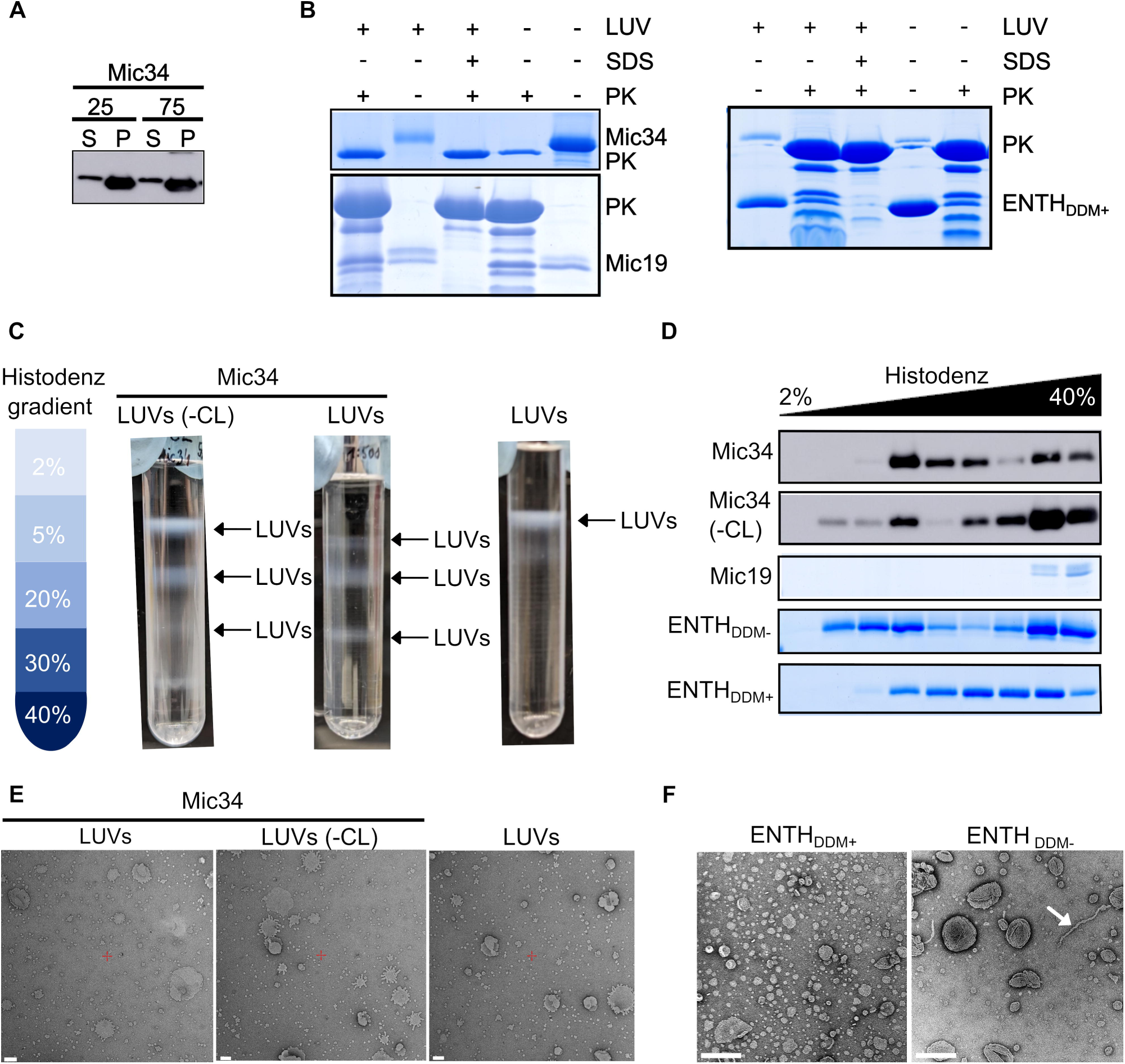
Mic34 binds to phospholipid bilayers *in vitro*. **(A)** Carbonate extraction of Mic34 loaded large unilamellar vesicles (LUVs) with 25 or 75 mM sodium carbonate. Supernatant, S; pellet, P. **(B)** Proteinase K (PK) protection assay of LUVs loaded with Mic34 or Mic19 and Folch liposomes loaded with ENTH visualized with Coomassie Brilliant Blue staining after SDS-PAGE. **(C)** Flotated LUVs with and without (-CL) cardiolipin separated on aa noncontinuous Histodenz density gradient (schema on left) loaded with recombinant Mic34. Compare banding pattern with LUVs flotated alone (right). **(D)** Histodenz density gradient fractions resolved by SDS-PAGE and stained with Coomassie Brilliant Blue or transferred to a membrane for incubation with an anti-His antibody (Mic34 with and without CL). **(E)** Negatively stained LUVs loaded with Mic34. Scale bar, 100 nm. **(F)** Negatively stained Folch + 5% PI(4,5)P2 liposomes incubated with the membrane remodeling protein ENTH that did (+DDM) or did not (DDM-) undergo prior solubilization with 0.1% DDM. Scale bar, 200 nm.

Next, we assayed whether rMic34 deformed the LUVs by flotation on Histodenz density gradients (Fig. 3C). As previously reported (Barbot et al. 2015; Tarasenko et al. 2017), LUVs alone float to the lightest fractions of the gradient. After incubation with rMic34, LUVs with or without cardiolipin formed multiple bands in denser fractions of the gradient, reminiscent of canonical Mic60 (Tarasenko et al. 2017). The presence of rMic34 in these fractions was verified by immunoblotting (Fig. 3D), indicating that rMic34 binding is weighing down the LUVs. The signal in the heaviest fraction corresponds to unbound rMic34, as seen in the rMic19-incubated LUV negative control.

To see if the striated banding pattern we observed often corresponds to LUV deformation (Barbot et al. 2015; Tarasenko et al. 2017), LUVs that flotated into lower density fractions were isolated and observed by negative-stain transmission electron microscopy (TEM). However, we could not reliably observe any obvious differences between the naked LUVs and those incubated with rMic34, independently of the presence of cardiolipin in the LUV membrane (Fig. 3E).

This lack of LUV deformation may be because rMic34 does not have *in vitro* membrane deformation activity or this activity is inhibited by the 0.1% DDM treatment. To test the latter hypothesis, we compared the liposome deformation activity of Epsin N-terminal Homology Domain (ENTH) that underwent 0.1% DDM solubilization prior to incubation with Folch liposomes spiked with 5% PI(4,5)P2 (DDM+) to that of untreated ENTH (Gleisner et al. 2016) (DDM-). DDM+ ENTH bound to the outside of liposomes (Fig. 3B-C). However, liposome tubulation was only observed in the absence of the detergent treatment (Fig. 3F). Thus, we are unable to address Mic34’s putative membrane remodeling activity *in vitro* with our current experimental set up. Nevertheless, we conclude that Mic34 binds phospholipids *in vitro* in a cardiolipin-autonomous way like yeast Mic60 (Hessenberger et al. 2017).

### Mic34 and Mic40 are capable of remodeling the cytoplasmic membrane of a gammaproteobacterium upon export to the periplasm

After establishing that Mic34 is able to bind phospholipid membranes, we next investigated whether Mic34 and Mic40 have membrane-bending activity *in vivo*. We employed an assay that has been used to demonstrate that Mic60 has this capacity (Tarasenko et al. 2017). Mic34 and Mic40 from *T. brucei* were heterologously expressed in the gammaproteobacterium *E. coli*. To facilitate their export into the periplasm, which represents the proteobacterial analog of the mitochondrial IMS, each MICOS subunit was fused to maltose binding protein (MBP).

First, *E. coli* transformed with various MBP expression constructs were separated into periplasm and spheroplast fractions (Fig. 4A). As expected, MBP alone was completely exported into the periplasm. This and the retention of T7 polymerase within the spheroplasts (supplementary fig. S7A, Supplementary Material online) verified successful fractionation. While a minor portion of Mic34-MBP and Mic40-MBP was observed in the periplasm, they were enriched in the spheroplast fraction (Fig. 4A). Carbonate extraction of the spheroplasts was done to see whether Mic34-MBP and Mic40-MBP were embedded in the membrane (Fig. 4A), like the integral subunit Mic24 fused to MBP (supplementary fig. S7B, Supplementary Material online). While some Mic34-MBP and Mic40-MBP were observed in the integral fraction–reminiscent of Mic34’s behavior in LUVs (Fig. 3A)–most accumulated in the soluble fraction (Fig. 4A). Our interpretation is that Mic34-MBP and Mic40-MBP tightly associate with periplasm-facing part of the cytoplasmic membrane because (1) the protein is partially observed in the periplasm fraction and (2) Mic34 and Mic40 are sorted into inclusion bodies when expressed in the cytoplasm (supplementary fig. S6A-B, Supplementary Material online).

**Fig. 4.**
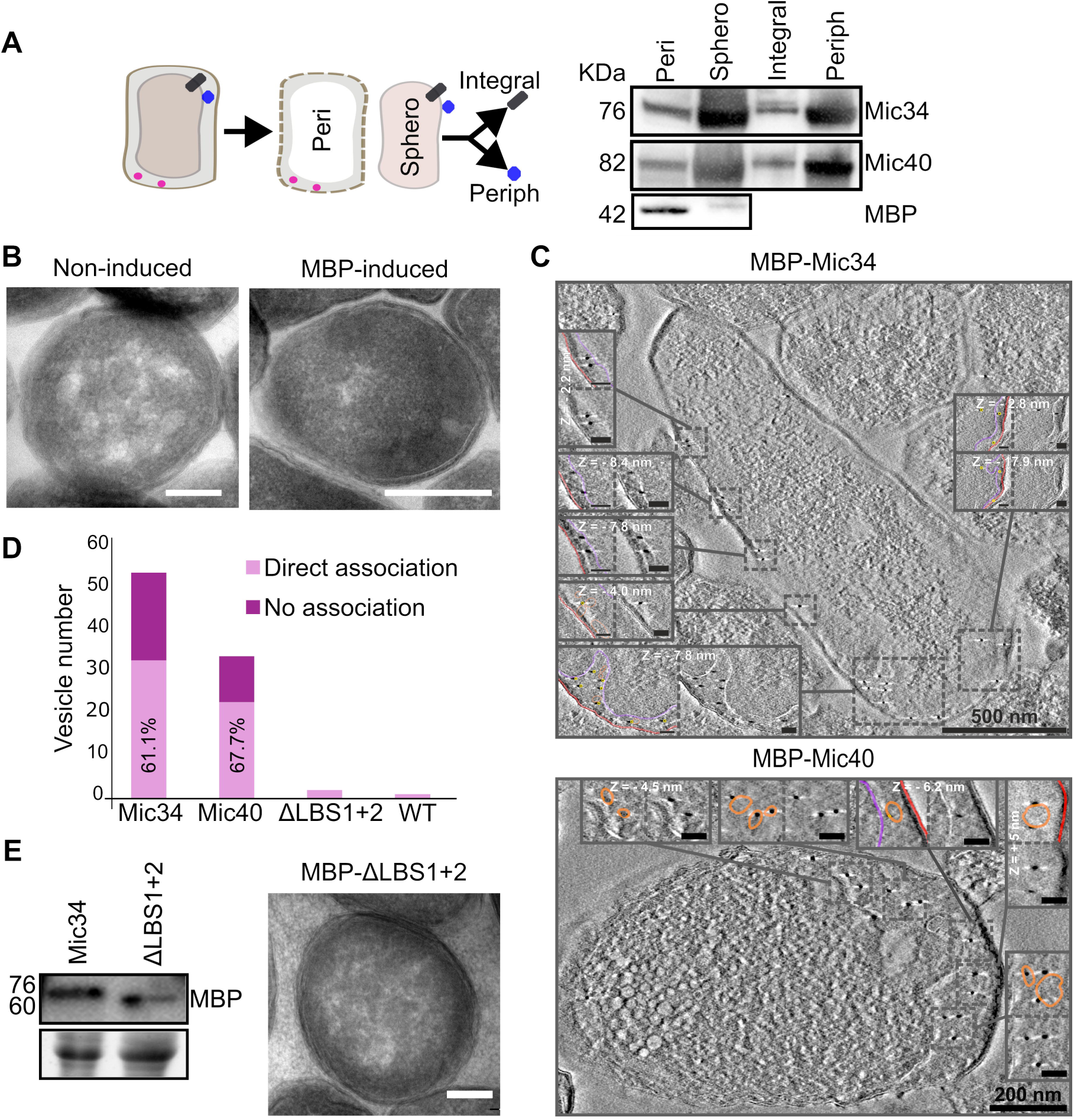
Heterologously expressed Mic34 and Mic40 remodels the cytoplasmic membrane of *E. coli* in an LBS-dependent manner upon periplasm export. **(A)** Schema (left) of periplasting by osmotic shock to separate spheroplast (*i.e. the* compartment bound by the cytoplasmic membrane) and periplasm fractions. The former was fractioned by carbonate extraction to separate integral and peripheral membrane proteins. Western blots on right show the localization of MBP-Mic34, MBP-Mic40, and MBP alone in *E. coli*, as detected with an anti-MBP antibody. **(B)** Representative transmission electron micrographs of *E. coli* when MBP expression is not (left) and is (right) induced. Scale bar, 200 µm. **(C)** Representative electron tomograms of immunogold-labeled MBP-Mic34 and MBP-Mic40 using an anti-MBP antibody. Full cells are shown in the main image with dashed boxes corresponding to magnified inset images that depict either unmarked or traced images. In the latter, pink line traces the cytoplasmic membrane, red the outer membrane, and orange the vesicles; yellow circles highlight electron-dense gold nanoparticles. Scale bars: main image, indicated above bar; inset, 50 nm. Insets report the depth in the Z-dimension of the depicted plane of the tomogram when applicable. **(D)** Bar graph reporting the total number of vesicles observed in TEM images (n=22) of cells expressing MBP tagged Mic34, Mic40, Mic34ΔLBS1+2, and of WT cells. The number of vesicles arising in Mic34 and Mic40 expressing cells. Vesicles directly associated with nanoparticles are shaded in light purple with their proportion shown. **(E)** Western blot probed using an anti-MBP antibody to show MBP-Mic34 and MBP-Mic34ΔLBS1+2 expression (left) and a representative TEM image of *E. coli* expressing MBP-Mic34ΔLBS1+2. Scale bar= 200 nm).

Next, we looked if Mic34 and Mic40 remodeled *E. coli* membranes similarly to heterologously expressed full-length *S. cerevisiae* Mic60 or just its IMS part (Tarasenko et al. 2017). After establishing by TEM that we can replicate this experiment (supplementary fig. S7C, Supplementary Material online), including reproducing that MBP alone does not affect *E. coli* ultrastructure (Fig. 4B), we observed Mic34-MBP and Mic40-MBP expressing *E. coli* by electron tomography (Fig. 4C) and TEM (supplementary fig. S5D, Supplementary Material online). Mic34-MBP and Mic40-MBP had respectively 5 and 3 times more vesicles observed by TEM in the periphery of the cell as compared to the parental strain (Fig. 4D). Immunogold labelling of the MBP moiety showed that both chimeras were also observed in the periphery, associated with the majority of vesicles. Electron tomography allowed us to better resolve these vesicles, which accumulated in the periplasm (Fig. 4C-D; supplementary movies S1 and S2, Supplementary Material online). Furthermore, we used this method to determine that the bulk of MBP-visualizing gold nanoparticles were associated with the vesicles (supplementary fig. S7E, Supplementary Material online). Together these results indicate that Mic34-MBP and Mic40-MBP are directly responsible for cytoplasmic membrane remodeling when heterologously expressed in *E. coli*, phenocopying *S. cerevisiae* Mic60 in this assay.

To test whether the predicted LBS helices of the slightly more conventional Mic34 (Fig. 1; supplementary fig. S7F, Supplementary Material online) are important for membrane remodeling, as has been demonstrated for *S. cerevisiae* Mic60 (Hessenberger et al. 2017), we expressed a periplasm-targeted LBS deletion mutant (Mic34ΔLBS1+2). We first tested whether the ∼6 kDa smaller mutant was expressed (Fig. 4E) and targeted to the periplasmic face of the *E. coli* cytoplasmic membrane (supplementary fig. S7A, Supplementary Material online). As expected, this mutant failed to remodel the cytoplasmic membrane, having about the same number of vesicles as the parental strain (Fig. 4D-E).

Wondering whether Mic34 and Mic40 can simultaneously remodel the cytoplasmic membrane, we transformed *E. coli* with two plasmids respectively encoding Mic34-MBP or Mic40-MBP. Co-expression was observed by gel staining, which revealed strong bands corresponding to the theoretical molecular weights of each chimera (supplementary fig. S8A, Supplementary Material online). This result was verified by immunoblotting with antibodies specifically recognizing Mic34 and Mic40. These cells exhibited prominent vesicles within the cytosol that were absent from the negative control when observed by electron tomography (supplementary fig. S8B-F, Supplementary Material online). This result suggests that Mic34 and Mic40 may work in concert, evocative of how Mic60 self-interaction mediates membrane remodeling activity (Bock-Bierbaum et al. 2022).

In conclusion, with an assay based on heterologous expression in *E. coli*, we were able to show that (1) Mic34 and Mic40 have the capacity to remodel membranes, and (2) they may act together to do so, and (3) Mic34’s LBS is directly involved in this activity. Thus, we provide empirical evidence that Mic34 and Mic40 are extremely diverged mitofilin family proteins.

### Overexpression of Mic34 causes reticulation of a morphologically simplified trypanosomal mitochondrion

The long slender bloodstream form (BSF) *T. brucei* has a morphologically simplified mitochondrion that runs as a tube along the cell body opposite the flagellum, reflecting the cessation of cellular respiration in this life cycle stage (Jakob et al. 2016; Hughes et al. 2017; Bílý et al. 2021). All MICOS complex subunits are downregulated in this stage compared to PCF (Kaurov et al. 2018). Thus, expression of electron transport chain subunits is necessarily triggered during BSF differentiation into PCF (Zíková 2022), corresponding with mitochondrial reticulation.

Ectopic expression of V5-epitope tagged Mic34 (Mic34-V5^+^) with 1000 ng/ml doxycycline in wild type BSF was considerably higher than that of endogenous Mic34 (Fig. 5A). Surprisingly, this overexpression reproducibly induced cytostasis after 24 hours, with induction by even 10 ng/ml doxycycline effectively inhibiting growth (Fig. 5A; supplementary fig. S9A-B, Supplementary Material online). Mic40-V5^+^ ectopic expression only mildly inhibited growth (supplementary fig. S9B, Supplementary Material online), perhaps due to its lower expression (supplementary fig. S9A, Supplementary Material online). After verifying that Mic34-V5^+^is targeted to the mitochondrion (supplementary fig. S9C, Supplementary Material online), we assayed how mitochondrial morphology was affected after 36 hours of Mic34-V5^+^ due to its cytostatic effect. Many Mic34-V5^+^ cells exhibited profound reticulation of their mitochondrion (Fig. 5C), with ∼2.3 times more branching than in the wild type BSF organelle (Fig. 5D). Apparent fragmentation of the organelle was also observed (supplementary fig. S9D, Supplementary Material online). However, we did not notice any obvious changes to the cristae that were sporadically observed by TEM (supplementary fig. S9E, Supplementary Material online). This is consistent with the uninterrupted absence of electron transport chain subunits and unaffected F_O_F_1_-ATP synthase expression in Mic34-V5^+^ cells (supplementary fig. S9F, Supplementary Material online). Furthermore, IMS protein levels were not altered, unlike during Mic34 depletion in PCF (Eichenberger et al. 2019). Thus, the observed morphological changes are likely due to Mic34-V5^+^ acting on the mitochondrion directly.

**Fig. 5.**
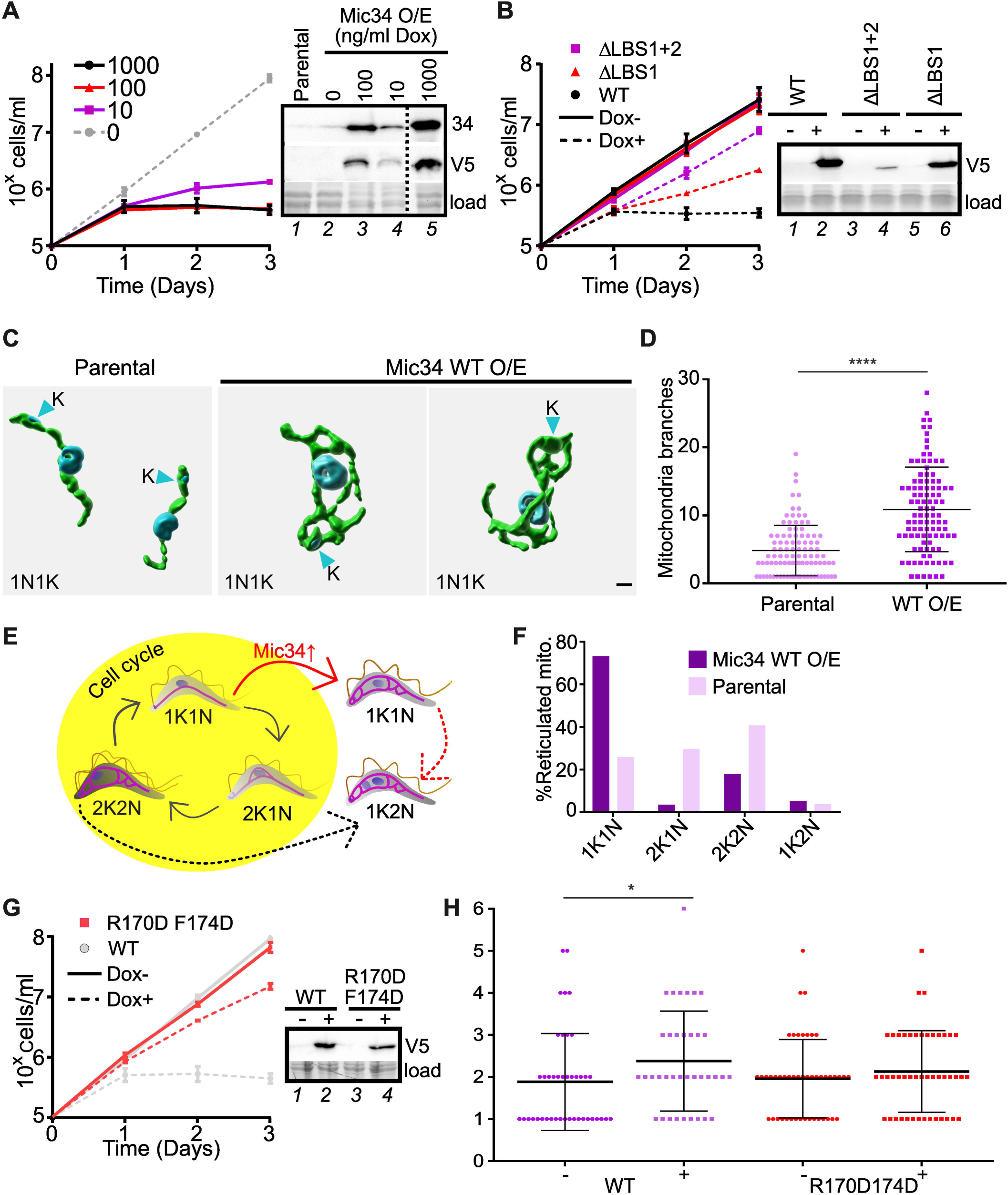
Mic34 overexpression induces reticulation of the tubular mitochondrion of bloodstream form *T. brucei*. **(A)** Growth of Mic34 overexpression (O/E) as measured by cell density plotted logarithmically under different doxycycline (Dox) concentrations (left). Solid line with circles, 1000 ng/ml; triangle,100 ng/mL; square,10 ng/ml. Dashed line, 0 ng/ml Dox. Standard deviation at each time point (n = 3) is not visible at this scale. Right, western blots probed with anti-Mic34 (detecting both endogenous Mic34 and V5-tagged Mic34) and anti-V5 tag antibodies, with fluorescently detected proteins used as a loading control (load). **(B)** Left, growth of cells overexpressing Mic34 WT (black circle), ΔLBS1+2 (purple square), ΔLBS1 (red triangle) by induction with 1000 ng/ml Dox (dotted line) compared to uninduced cells (solid lines). Standard deviation at each time point (n = 3) is not visible at this scale. Right, western blot probed with anti-V5 to show expression of Mic34 WT, ΔLBS1+2, and ΔLBS1 with (+) and without (–) 100 ng/ml Dox induction. **(C)** Representative 3D reconstructions of mitochondria in parental and WT Mic34 O/E cells. Mitochondria labeled by anti-HSP70 antibody are in green; the DAPI-stained nucleus and kinetoplast DNA (arrowhead) are in blue. Scale bar, 1 µm. Note all cells are 1N1K cells. **(D)** Quantification of mitochondrial branching in 100 parental and Mic34 WT O/E (36 hours after Dox induction) cells. Statistical significance: ****, p < 0.0001. **(E)** Schema of the *T. brucei* bloodstream cell cycle under normal conditions (yellow background with black arrows) and upon Dox-induced Mic34 overexpression (↑Mic34 O/E) (red arrow). Mitochondria are in blue and the number of nuclei (N) and kinetoplast DNA (K) is given below each cell. Dashed arrows indicate a pathway to defective 1N2K cells under normal Mic34 expression and overexpression. **(F)** Bar graph showing the percentage of cells with reticulated mitochondria from different cell cycle stages depicted in E in parental and WT Mic34 overexpression cells. **(G)** Growth of *T. brucei* overexpressing Mic34^R170DF174D^ with 1 µg/ml of doxycycline (Dox+) compared to uninduced (Dox-) cell lines. Standard deviation whiskers at each time point (n = 3) are faintly visible at 3 days in this scale. This overlays the growth 1 µg/ml of doxycycline-induced Mic34 WT and its uninduced counterpart from A (grey). Right, western blots probed with anti-V5 tag antibody (top) and loading control (bottom; load) as in A. **(H)** Quantification of mitochondrial fragmentation in Mic34 WT and Mic34^R170DF174D^ overexpressing (+, 36 hours after Dox induction) and non-induced (-) cells. Statistical significance: *, p < 0.05. Number of cells counted: WT-, 42; WT+, 37; Mic34^R170DF174D^ (both + and -), 47.

We were concerned that Mic34-V5^+^ branching phenotype was an artifact of cell cycle arrest. The mitochondrion becomes more branched during cell cycle progression to ensure inheritance of the organelle after cytokinesis (Jakob et al. 2016; Hughes et al. 2017). Thus, a defect to this process could secondarily cause a reticulated mitochondrion (Fig. 5E). To address this issue, we counted the number of kinetoplasts and nuclei in Mic34-V5^+^ cells bearing a reticulated mitochondrion; branching starts during kinetoplast division, perceptively increasing as the cell goes from having a single nucleus and kinetoplast (1N1K) to two separate kinetoplasts (1N2K) and eventually two separate nuclei (2N2K) (Fig. 5E). The vast majority of unsynchronized Mic34-V5^+^ cells with a reticulated mitochondrion were 1N1K (Fig. 5C, F), at early cell cycle stages that normally have a tube-like mitochondrion. As expected, wild type BSF exhibited branched mitochondria mostly in the later 2N2K cell cycle stage. Thus, we conclude that Mic34-V5^+^ overexpression directly affects mitochondrial morphology.

We hypothesized that the Mic34 LBS alpha helices may be important for the observed cell cycle arrest due to mitochondrial remodeling. We overexpressed mutants in which both (Mic34ΔLBS1+2^+^) or just LBS1 (Mic34ΔLBS1^+^) were deleted (supplementary fig. S7F, Supplementary Material online). Like WT Mic34, these mutants were also targeted to the mitochondrion (supplementary fig. S9C, Supplementary Material online). After 24 hours of overexpression growth inhibition correlated with the extent of LBS deletion (Fig. 5A). The mild growth inhibition in Mic34ΔLBS1+2^+^ could be explained by its relatively lower expression compared to others (Fig. 5B, *c.f.* lane 4 with 2 and 4). However, Mic34ΔLBS1+2^+^ and Mic34-V5^+^ induced with 10 ng/ml doxycycline (Fig. 5A, lane 4) are equivalent, with the latter exhibiting more dramatic growth inhibition than the deletion mutant. Together with our previous *E. coli* data, we conclude that the LBS of the mitofilin-like Mic34 plays an important functional role as it does in canonical Mic60.

Tertiary (Fig. 1B-C) and primary structural alignments predicted that the conserved LBS1 phenylalanine of *T. brucei* LBS1 (F175) would be a critical residue for the observed Mic34 mitochondria remodeling given it and an upstream arginine (R) residue were shown to be essential for yeast Mic60 remodeling activity (Hessenberger et al. 2017). We expressed a double point mutant affecting these amino Mic34^R170D_F174D^, which was equivalently expressed compared to Mic34-V5+)(Fig 5G; supplementary fig. S9C, Supplementary Material online) and targeted to the mitochondrion (supplementary fig. S9C, Supplementary Material online). The growth of Mic34^R170D_F174D^ was restored to almost the same rate as that of the uninduced counterpart (Fig. 5G), which may be explained by a mitigation of apparent fragmentation seen in Mic34^R170D_F174D^ compared to Mic34-V5^+^ (Fig. 5H; supplementary S9D, Supplementary Material online). Thus, these conserved amino acid residues are essential for the dominant negative effect seen upon expression of Mic34-V5^+^. We conclude that Mic34 was directly responsible for the mitochondrial remodelling observed upon its overexpression. Furthermore, the cytostatic effect of Mic34-V5^+^ may be linked with the apparent fragmentation that also occurs.

## DISCUSSION

Here, we provide *in silico* and experimental evidence supporting the conclusion that euglenozoan MICOS contains two soluble subunits that represent cryptic mitofilin family proteins. In contrast to Mic60 from two domains of life, Mic34 and Mic40 do not contain TMD that embeds them into a phospholipid bilayer, as supported by its repeated association with the periphery of membranes in this and another (Kaurov et al. 2018) study. It has been proposed before in passing (Hashimi 2019; Prokopchuk et al. 2023) that the ancestral Mic60 got split into two parts, one represented by the integral membrane Mic24 and the other by the soluble Mic34 in the forebear of Euglenozoa. Our analyses did not provide any evidence that Mic24 evolved from any part of the ancestral Mic60 gene, and its origin remains obscure. However, our results corroborate that Mic34 indeed represents a cryptic mitofilin domain-containing protein. Additionally, we demonstrate that the still hypothetical split of integral and soluble components was followed by duplication of the latter, mitofilin domain-containing part, resulting in the paralogous Mic34 and Mic40 pair. Admittedly, the phylogenetic tree inferred from mitofilin domain sequences is not well resolved and does not support this scenario, as it would need Mic34 and Mic40 to branch most closely to Mic60 from Heterolobosea (the sister group to Euglenozoa). However, this is not unexpected, given the sequence divergence of the sampled mitofilin family proteins and the resulting challenges for phylogenetic inference; note also that the branching order of Mic60 itself conforms less to organismal phylogeny than that of Mic34 and Mic40.

The presence of cryptic mitofilin domains in Mic34 and Mic40 predicts that these are membrane remodeling proteins like conventional Mic60 from prokaryotes and eukaryotes. However, we were unable to verify Mic34 and Mic40’s capacity to remodel phospholipid bilayers *in vitro* here. Nevertheless, this assay did reveal Mic34 can bind liposomes. Thus, we had to resort to assaying their putative role in membrane remodeling by other means. Heterologous expression of Mic34 and Mic40 deformed *E. coli* cytosolic membranes, the prokaryotic analog to the mitochondrial IM (Muñoz-Gomez et al. 2017; Muñoz-Gómez et al. 2023), as yeast and alphaproteobacterial Mic60 did (Tarasenko et al. 2017). However, a difference should be noted here: Mic60 caused the cytoplasmic membrane to invaginate inward into the cytoplasm whereas Mic34 and Mic40 promoted the accumulation of vesicles within the IMS. Furthermore, Mic34 and Mic40 may work together as their co-expression in *E. coli* results in conspicuous membrane remodeling. This suggests that Mic34 and Mic40 interact with each other *in vivo*, as conventional Mic60 acts on membranes as a homooligomer (Bock-Bierbaum et al. 2022). Mic34 requires the LBS motifs it shares with Mic60 (Hessenberger et al. 2017) for cytoplasmic membrane vesicularization, consistent with the notion it is a cryptic member of the mitofilin protein family.

Mic34 ectopic expression in a *T. brucei* BSF with a reduced, tubular mitochondrion causes elaboration into a branched organelle. This parallels the enhanced interconnectivity of mitochondria upon Mic60 overexpression in primary and immortalized neuronal cell lines (Van Laar et al. 2016). These assayed Mic34 membrane-deformation activities *in vivo* rely on its predicted LBS motifs, further supporting Mic34 and its paralog Mic40 as diverged mitofilin protein family members. LBS1 contains a conserved aromatic amino acid (Muñoz-Gómez et al. 2023) that is essential for Mic60’s *in vitro* and *in vivo* activities (Hessenberger et al. 2017). This residue along with an upstream arginine residue is also essential for Mic34-mediated BSF mitochondria remodeling, as predicted. The aromatic residue in *C. thermophila* Mic60 is buried inside the hydrophobic interaction interface of the Mic60 homodimer (Bock-Bierbaum et al. 2022). Whether this residue plays a similar role in Mic34 homooligomerization and/or heterooligomerization remains an open question.

Mic34 overexpression did not alter cristae dramatically in BSF *T. brucei*, contradicting Mic60 overexpression phenotypes in yeast (Rabl et al. 2009) and alphaproteobacteria (Muñoz-Gómez et al. 2023), manifested in the branching of their respective bioenergetic subcompartments (*e.g.* cristae). However, this inconsistency can be explained by the lack of an active electron transport chain in BSF (Bílý et al. 2021; Zíková 2022), which were present when Mic60 was overexpressed in yeast and bacteria. Thus, an electron transport chain may be necessary for populating and stabilizing elaborated crista membranes. Alternatively and/or additionally, co-expression in *E. coli* implies Mic34 interacts stoichiometrically with Mic40 and/or (an)other MICOS subunit(s) to shape crista.

One clear divergence from canonical Mic60 is that Mic34 (Eichenberger et al. 2019) and Mic40 (this study) appear to be directly integrated into the MIA pathway for IMS protein import in PCF, like the thioredoxin-like Mic20 subunit (Kaurov et al. 2018; Kaurov et al. 2022). In contrast, while yeast Mic60 augments the Mia40 sulfhydryl oxidase’s mediation of MIA (von der Malsburg et al. 2011), Mic60 deletion does not lead to depletion of MIA substrates (Varabyova et al. 2013) like Mic34 and Mic40 silencing. The MIA defects seen upon depletion of Mic34 and Mic40 complicate the interpretation of crista defects given that putative respiratory chain subunits and assembly factors, plus the small Tim proteins, are imported via MIA. Moreover, the latter are needed for import of IM proteins (Wenger et al. 2017; Smith et al. 2018), many of which are assembled into respiratory chain complexes (Zíková 2022). Based on our results, the MIA pathway also is important for maintaining reticulated mitochondria in PCF, as shown by Erv1 simplifying mitochondrial complexity to the same degree as Mic34 and Mic40. However, it is possible that concomitant downregulation of the soluble MICOS subunits Mic20 and Mic32–and by extension, perhaps the rest of this subcomplex–may be responsible for the morphological phenotype of Erv1 RNAi after all (Peikert et al. 2017). Thus, at this time we cannot verify nor refute the hypothesis that MICOS plays a role in maintaining a reticulated mitochondrial network *via* its demonstrated membrane remodeling activity.

It is worth noting that this work has implications about the subcomplex architecture of MICOS throughout eukaryotes. In opisthokonts, two adjacent integral membrane subcomplexes are each assembled around their respective core subunit, Mic10 and Mic60 (Hashimi 2019; Colina-Tenorio et al. 2020). It is reasonable to expect this ‘horizontal’ (relative to the membrane) configuration in the majority of aerobic eukaryotes because they have both Mic10 and Mic60 (Muñoz-Gómez et al. 2015; Huynen et al. 2016). Uniquely, trypanosomal–and by extension likely also euglenazoan–MICOS has a vertical architecture: a membrane-embedded subcomplex containing two Mic10 paralogs and a peripheral subcomplex comprised of soluble subunits located within the IMS (Eichenberger et al. 2019). This divergent subcomplex architecture likely corresponds to the soluble nature of IMS localized Mic34 and Mic40 (Kaurov et al. 2018). Significantly, the existence of the Mic34/Mic40 subcomplex implies that a mitofilin domain represents the core of a MICOS module regardless of whether or not it is directly IM anchored. Perhaps the Mic60 subcomplex is more predisposed to IM detachment than the Mic10 subcomplex. In opisthokonts, the only soluble MICOS subunit Mic19 (plus its paralog Mic25 in vertebrates) is part of the Mic60 complex (Muñoz-Gómez et al. 2015; Huynen et al. 2016; Hessenberger et al. 2017; Bock-Bierbaum et al. 2022). In contrast, the persistence of the trypanosomal Mic10 subcomplex within the IM suggests it is perpetually an integral IM MICOS module.

In the diverged euglenozoans and opisthokonts, a mitofilin protein represents the largest subunit in the MICOS complex. This may reflect the mitofilin domain’s role in bridging juxtaposed membranes at contact sites and crista junctions as well as being an interaction hub to facilitate coordination of MICOS with other fundamental pathways (Aaltonen et al. 2016; Michaud et al. 2016). Such a central role in MICOS is likely a product of the mitofilin domain’s ancient, pre-mitochondrial origin (Muñoz-Gómez et al. 2023). Unlike the rather widespread loss of Mic60 in anaerobes without cristae, the transfiguration of Mic60 into Mic34/Mic40 appears to be a singular event that occurred exclusively in Euglenozoa, joining other unique mitochondrial apomorphies of this phylum (Cavalier-Smith 2016).

Our work here also shows that the mitofilin protein family is more diverse than originally thought, given the conservation of Mic60 in alphaproteobacteria and mitochondria (Muñoz-Gómez et al. 2023). It is unknown whether the duplication of the soluble mitofilin proteins is an adaptation to loss of their respective TMDs, or perhaps the divergence of Mic34 and Mic40’s mitofilin domains necessitates loss of their membrane integration. Perhaps the duplication of the mitofilin domain, resulting in the Mic34 and Mic40 paralogs during early euglenozoan evolution, allows one of the mitofilin domains to be fine-tuned to the needs of the subdivisions Euglenida, Diplonemea, and Kinetoplastea. This reveals surprising variability of the mitofilin family of proteins found in the organismal diversity occurring in two domains of life.

## MATERIAL AND METHODS

### Gathering of euglenozoan protein sequence datasets

Homologs of *T. brucei* Mic34 and Mic40 were identified by a combination of blast (Altschul et al. 1997) and HMMER3 searches (Eddy 2011) in a variety of databases and sequence resources, including the NCBI non-redundant protein sequences database, TriTryp (Alvarez-Jarreta et al. 2024), EukProt v3 (Richter et al. 2022), and sequence assemblies and annotations accompanying individual publications (details provided in supplementary dataset S1, Supplementary Material online). Briefly, homologs were first collected with blast (considering hits with e-value <1e-5) and aligned with MAFFT v7 (https://mafft.cbrc.jp/alignment/server/; (Katoh et al. 2019)). (Katoh et al. 2019)The HMMER3 package was then used for building a profile HMM for each MSA and searching the target protein sequence sets (using the hmmbuild and hmmsearch command, respectively). Newly identified hits missing in the original MSA and passing the program-defined inclusion threshold were collected, added to the MSA and the whole search was repeated until no new significant hits were retrieved or the search started to retrieve hits that were found not to represent euglenozoan Mic34 or Mic40 proteins. In a few cases homologs were obtained by tblastn searches against genome or transcriptome assemblies and the sequence of the respective encoded proteins were deduced by conceptual translation.

Only a subset of identified sequences was retained for further analyses, taking into account the degree of their completeness, whether both Mic34 and Mic40 homologs were identified for the given species (some of the sequences resources, especially the various euglenid single-cell transcriptomes, are very incomplete and did not yield both expected homologs), and keeping the taxon sampling balanced (i.e. avoiding the inclusion of highly similar sequences from closely related species). As a result, we obtained a set of Mic34 and Mic40 homologs from 44 euglenozoan taxa encompassing representatives of all three main euglenozoan clades as well as their various sublineages, In a few cases the sequences were attributed to a wrong organism in the given database, stemming from contamination of the source protist cultures used for transcriptome sequences by a co-cultured euglenozoan, whose identity was determined by investigating informative marker sequences present in the same assembly, such as the 18S rRNA gene or elongation factor 2 (EF2) protein. All the sequences were manually curated to achieve their accuracy (e.g. by removing artificial N-terminal extensions frequently present in protein sequences automatically inferred from transcriptome assemblies); in some cases the existing truncated sequences were extended or even completed by manually recruiting original Illumina sequencing reads identified by iterative blast searches against the respective sequencing runs in the Sequence Read Archive (SRA; https://www.ncbi.nlm.nih.gov/sra).

Homologs of the *T. brucei* Mic24 protein (Tb927.9.10160) were searched and curated in a manner analogous to that used for gathering the Mic34 and Mic40 sequence datasets. The euglenid homologs retrieved with a profile HMM derived from a MSA of kinetoplastid Mic24 proteins included a protein from *Euglena gracilis* that was listed as a putative ortholog of *T. brucei* Mic24 (=Mic60) in a previous study (Hammond et al. 2020). No candidates for Mic24 homologs were identified in any diplonemid and *Perkinsela* spp. even using sensitive HMMER searches queried with profile HMMs of kinetoplastid, euglenid, or combined kinetoplastid-euglenid Mic24 for MSAs, suggesting loss or extreme divergence of Mic24 in these lineages. We likewise failed to identify a Mic24 homolog in the euglenid *Ploeotila vitrea* (a species included in our selection of taxa for the Mic34 and Mic40 analysis), but incompleteness of the available sequence data (a transcriptome assembly) cannot be excluded in this case. All the Mic34, Mic40, and Mic24 sequences included in the final datasets and the related metadata are provided in supplementary dataset S1, Supplementary Material online.

### HHpred searches

Homology of Mic34 and Mic40 to other proteins was rigorously examined with HHpred (Söding et al. 2005) as available at the MPI Bioinformatics Toolkit server (https://toolkit.tuebingen.mpg.de/tools/hhpred). The searches with queries representing individual sequences, i.e. *T. brucei* Mic34 and Mic40, were done with the default parameters, i.e. with initial three iterative HHblits searches against the UniRef30 database, which in both cases retrieved a set of exclusively kinetoplastid homologs that were then included by the program into the query MSA, used then for building a profile HMM and searching the target database of profile HMMs based on HMM-HMMM comparisons. In the case of searches in which the query was the custom Mic34 or Mic40 MSA (in each case computed with MAFFT from the final set of 44 selected euglenozoan representatives), no initial HHblits searches were allowed and the query MSA was directly used for creating the query profile HMM. For reference we report results of HHpred searches against the Pfam-A_v37 database, which all retrieved the Pfam database entry PF09731.14 as the only significant hit. Alternatively, we also searched the COG_KOG_v.1 database and identified with both Mic34 and Mic40 (individually or as an MSA) the same entry as the best (and significant) hit, i.e. KOG1854, which represent eukaryotic Mic60 proteins, reinforcing the results of the searches against Pfam-A. The relative divergence of the mitofilin domains of the euglenozoan Mic34 and Mic40 proteins and of Mic60 proteins from other eukaryotes was evaluated by carrying pairwise profile HMM-HMM comparisons with HHpred, using as the query and the target the respective custom MSAs (the same as used in the phylogenetic analysis; see the next section).

### Phylogenetic analysis of the mitofilin domain

A set of reference eukaryotic Mic60 sequences was gathered by selecting proteins from a broader sequence set obtained by searching the EukProt v3 database with HMMER3 and a profile HMM built from the seed alignment of the Pfam family PF09731 (mitofilin). Sequences were selected to include a representative from different major eukaryote lineages, with a denser sampling for the supergroup Discoba, owing to the presumed origin of the euglenozoan Mic34 and Mic40 from an ancestral discoban Mic60. Additional discoban Mic60 sequences were identified by HMMER or blast searches in relevant sequence resources from species not included in EukProt v3 and added to the dataset (details provided in supplementary dataset S1, Supplementary Material online). The Mic60 sequences were combined with the Mic34 and Mic40 sequences used in HHpred analyses (see above) and a multiple alignment was built using MAFFT v7 and the L-INS-I iterative refinement method. The alignment was restricted to the C-terminal region corresponding to the mitofilin domain and used for tree inference performed the ETE3 3.1.3 pipeline (Huerta-Cepas et al. 2016) as implemented on the GenomeNet server (https://www.genome.jp/tools-bin/ete). Specifically, the alignment was trimmed with). Specifically, the alignment was trimmed with trimAl v1.4.rev6 (Capella-Gutiérrez et al. 2009), removing columns with >20% of gaps, and the tree was inferred using RAxML v8.2.11 (Stamatakis 2014), with the default parameters, the substitution model PROTGAMMAJTT, and 100 bootstrap replicates. The resulting tree was visualized in an arbitrarily rooted form using iTOL v6 (Letunic and Bork 2024) and further processed using a graphical editor. v1.4.rev6 (Capella-Gutiérrez et al. 2009), removing columns with >20% of gaps, and the tree was inferred using RAxML v8.2.11 (Stamatakis 2014), with the default parameters, the substitution model PROTGAMMAJTT, and 100 bootstrap replicates. The resulting tree was visualized in an arbitrarily rooted form using iTOL v6 (Letunic and Bork 2024) and further processed using a graphical editor.

### Structural predictions

The HHpred server was used to detect the mitofilin domain of Mic34 and Mic40 using hidden Markov model profiles to interrogate the following manually set structural and domain databases: PDB_mmCIF70_18_Jun; Pfam-A_v36; UniProt-SwissProt-viral70_3_Nov_2021; TIGRFAMs_v15.0. E-values reported in Fig. 1A are for the top hit ‘PF09731.12; Mitofilin’. AlphaFold 2 predictions for *T. brucei* Mic34 and Mic40 and *N. gruberi* Mic60 were performed according to a previously established pipeline catered to discoban proteins (Wheeler 2021). Mitofilin domains were aligned in ChimeraX by restricting the matchmaker to the two domains (Meng et al. 2023). The mitofilin alpha helices were mapped by manual inspection of the aforementioned structural predictions and crosschecked with multiple sequence alignments to identify deletions and fusions in the mitofilin domains.

### *Trypanosoma brucei* culturing, transfection, fractionation and growth analysis

The PCF Mic34, Mic40 and Erv1 inducible RNAi and SMOXp parental (Poon et al. 2012) cell lines were generated with the same constructs plus strategy and then grown as previously described (Haindrich et al. 2017; Kaurov et al. 2018). The pleomorphic AnTat1.1 BSF cell line was cultivated in HMI-9 media (Hirumi and Hirumi 1989) at cell densities ≤8 × 10^5^ cells/ml to maintain pleomorphism. Cells were transfected using a Amaxa Nucleofector (X-001) with the pT7-3xV5 vector (Flaspohler John et al. 2010), in which wild type and mutated *mic34* and *mic40* was cloned to facilitate their inducible ectopic expression by addition of 1 µg/ml of doxycycline to the media. Cells were fractionated into cytoplasmic and organellar fractions as previously described (Kaurov et al. 2018). Cell densities were measured on a Beckham Coulter Z2 Cell and Particle Counter.

### Mass spectroscopy analysis

For depletome data, mitochondria from 10^8^ parental SMOXp, TP3 and TP5 Mic40i *T. brucei* cells were crudely isolated as before (Kaurov et al. 2018) and snap frozen in liquid nitrogen for storage at -80°C prior to LC-MS/MS mass spectroscopy analysis, which was performed and the data visualized by a previously established pipeline (Benz et al. 2022) with slight modifications. Briefly, for sample prep, SP4 (Solvent Precipitation SP3) protocol without beads was used (Johnston et al. 2022). Pellets of enriched mitochondria (50□μg of protein) were solubilized by SDS (final concentration 1% [w/v]), reduced with TCEP [tris(2-carboxyethyl)phosphine], alkylated with chloroacetamide and digested sequentially with Lys-C and trypsin. Samples were desalted on Empore C18 StageTips, dried in a speedvac, and dissolved in 0.1% TFA□+ 2% acetonitrile. About 0.5 μg of peptide digests were separated on a 50□cm C18 column using 90□min elution gradient and analyzed in DIA mode on an Orbitrap Exploris 480 (Thermo Fisher Scientific) mass spectrometer equipped with a FAIMS unit.

### Recombinant Mic34, Mic19 and ENTH expression and purification

The Mic34 open reading frame, omitting the region encoding the first 14 amino acid residues, was cloned into the pSKB3 vector via the BamHI and HindIII restriction sites in frame with the N-terminal His-tag sequence (Kaurov et al. 2022). The BL21(DE3)-derived C4 *E. coli* strain (Miroux and Walker 1996) was transformed with the plasmid. Upon reaching 0.6 OD_600_, rMic34 production was induced with 1 mM isopropyl β-D-1-thiogalactopyranoside (IPTG) for 3 h at 37°C. Isolated inclusion bodies containing rMic34 were dissolved in 8 M urea. This mixture was loaded into ÄKTAprime Plus liquid chromatography system connected the 1 ml HisTrap column (GE Healthcare), followed by washing with lysis buffer (8 M urea, 2 mM Tris pH 7.2, 200 mM NaCl, 1mM EDTA) at a 1 ml/min flow rate. An increasing gradient of elution buffer (lysis buffer with 400 mM imidazole) was introduced into the column; rMic34 was eluted at 40% (v/v) elution buffer. Fractions in which rMic34 was detected by Coomassie stained SDS-PAGE were pooled and loaded onto a HiLoad 16/600 Superdex 200 pg column (Cytiva) for size exclusion in 8 M urea, 150 mM NaCl, 10 mM Tris-HCl, pH 8. Purest fractions were chosen based on SDS-PAGE analysis and the protein was refolded through step-wise dialysis against refolding buffer (0.1% DDM, 150 mM NaCl, 20 mM HEPES, pH 7.4).

The *S. cerevisiae MIC19* coding sequence was cloned into the pGEX-6P-1 vector (Cytiva) via the BamHI and SalI restriction sites with a C-terminal TEV cleavage site and His-tag, and in frame with the N-terminal GST-tag. BL21(DE3) *E. coli* were transformed with the plasmid and grown to an OD600 of 0.6 before induction with 1 mM IPTG for 20 h at 18°C. Cells were lysed in lysis buffer (complete protease inhibitor cocktail without EDTA [Roche], 5% glycerol, 50 mM imidazole, 150 mM NaCl, 20 mM HEPES, pH 7.4), cleared by ultracentrifugation at 70000 × g for 25 min and loaded onto a His-Trap column (Cytiva) using an Äkta purifier chromatography system. Unbound proteins were washed out with 10 column volumes (CV) of lysis buffer and bound proteins eluted in 500 mM imidazole lysis buffer. Imidazole was dialysed out in dialysis buffer (5% glycerol, 150 mM NaCl, 20 mM HEPES, pH 7.4). The GST tag was removed via PreScission protease (Cytiva) and removal of GST and PreScission protease on glutathione Sepharose (GE Healthcare) and the His-tag via incubation with TEV protease in the presence of 14 mM β-mercaptoethanol and removal of His-tag and TEV protease on nickel-nitrilotriacetic acid (NiNTA) agarose beads (Macherey-Nagel).

The *Rattus norvegicus* ENTH domain of Epsin was purified based on a previously published protocol (Kroppen et al. 2021). In short, the protein was expressed with an N-terminal GST tag from the pGEX-6-P2 vector in BL21(DE3) cells. Expression was induced in LB medium with 1 mM IPTG for 3 h at 30°C. The cells were harvested and lysed in lysis buffer (complete protease inhibitor cocktail without EDTA [Roche], 150 mM NaCl, 50 mM HEPES, 1 mM EDTA, 1 mM DTT, pH 7.4). Lysed cleared by ultracentrifugation was loaded onto a GST-Trap column (Cytiva) using an Äkta purifier chromatography system. Bound protein was eluted in lysis buffer containing 10 mM reduced glutathione and peak fractions were concentrated. The GST tag was cleaved with Precision Protease at 4°C overnight. To separate cleaved GST, the buffer was then exchanged to low salt buffer (50 mM NaCl, 50 mM HEPES, pH 8) using a Sephadex G-25 column (GE Healthcare) and the sample applied onto a HiTrap Q column (Cytiva). The bound ENTH domain was eluted by steadily increasing the NaCl concentration to 1 M. Concentrated peak fractions were further purified by size exclusion on a HiLoad 16/600 Superdex 75 pg column (Cytiva) in 150 mM NaCl, 20 mM HEPES, pH 7.4, and ENTH-containing fractions were pooled.

### LUV preparation

Egg l-α-phosphatidylcholine (PC), egg l-α-phosphatidylethanolamine (PE), liver l-α-phosphatidylinositol (PI), brain l-α-phosphatidylserine (PS), Heart cardiolipin (CL), and brain L-α-phosphatidylinositol-4,5-bisphosphate (PI(4,5)P2) were purchased from Avanti Polar Lipids, Inc. Folch bovine brain extract Fraction I was purchased from Sigma Aldrich. IMM (48% PC/24%PE/16%PI/4%PS/8%CL), IMM-CL (56% PC/24%PE/16%PI/4%PS) and Folch + 5% PI(4,5)P2 lipid mixtures were dried under nitrogen flow to form thin lipid films. The IMM phospholipid mixture used to make LUVs was informed by Horvath and Daum (2013). Then they were dried in a desiccator for at least 3 h. Lipid films were hydrated in rehydration buffer (150 mM NaCl and 20 mM HEPES buffer, pH 7.4). Resulting LUVs were subjected to repeated freeze–thaw cycles and extruded through a 100-nm diameter polycarbonate filter (Whatman). Protein was added to liposomes in 0.1% DDM rehydration buffer with a molar protein/lipid ratio of 1:500.

### Flotation assay, sodium carbonate extraction, proteinase K protection

Flotation assays were performed in nonionic Histodenz (Sigma-Aldrich) density gradients. LUVs were placed on the bottom of the ultracentrifugation tubes, and a discontinuous 40/30/20/10/2% Histodenz density gradient was built up from high density to the lowest at the top of the tube. LUVs were ultracentrifuged for 1 h at 273,000 × g and 4°C. Afterwards, the gradient was dissected into nine equal fractions, precipitated with 10% TCA, and analyzed by SDS-PAGE. For sodium carbonate extraction and proteinase K protection assay, LUVs were flotated in a two-step density gradient consisting of 40% and 2% Histodenz. LUVs and co-flotating proteins were collected from the interphase. For sodium carbonate extraction, LUVs were incubated with 3 volumes of 100 mM Na_2_CO_3_, pH 11, for 30 min at RT before ultracentrifugation at 167,000 × g for 30 min at 4°C to separate membrane-bound from membrane-dissociated proteins. The supernatant containing lose protein was TCA-precipitated before both fractions were analysed by SDS-PAGE. For proteinase K protection assay, collected LUVs were washed in 8 volumes of rehydration buffer (150 mM NaCl, 20 mM HEPES, pH 7.4), pelleted at 167,000 × g for 30 min at 4°C and again resuspended in rehydration buffer before being incubated with 40 µg/ml proteinase K, or buffer, in the presence or absence of 1% SDS for 15 min at RT. Then, 2 mM PMSF was added and the samples incubated for 10 min at RT before being TCA-precipitated, resuspended in SDS-loading dye containing 1 mM PMSF and analysed by SDS-PAGE. As a control, proteins in absence of LUVs were treated in the same way.

### *Escherichia coli* culturing, transformation and subcellular fractionation

Coding sequences (CDSs) of *T. brucei* Mic34, Mic40 and Mic24 (omitting the regions encoding the first 14, 28, and 7 amino acid residues, respectively) were cloned into the BamHI and HindIII restriction sites of the pMAL-P2X vector and transformed into BL21(DE3) *E. coli*. The induction of MBP chimeras, which are targeted to the periplasm, was done as previously (Tarasenko et al. 2017), except using LB broth. The subcellular fractionation of *E. coli* was performed as before (Tarasenko et al. 2017), except the integral proteins remaining in the cytoplasmic membrane of the spheroplasts were separated by 0.1M Na_2_CO_3_ pH 11.5 treatment like previously (Kaurov et al. 2018).

For co-expression, the same Mic40 CDS was cloned into pMAL-C2X, which bears kanamycin resistance. BL21(DE3) *E. coli* were transformed concurrently with pMAL-P2X-Mic34 and pMAL-C2X-Mic40 and *E. coli* hosting both were selected in LB supplemented with 50 µg/ml kanamycin and 100 µg/ml ampicillin. *E.coli* lysates in were resolved on Mini-PROTEAN TGX Stain-Free Precast gels, which allow visualization of abundant proteins by UV illumination. Dual expression was verified by immunoblotting with antibody recognizing Mic34 (Eichenberger et al. 2019) and Mic40. The latter rabbit antibody was produced by Davids Biotechnologie GmbH (Regensburg, Germany) using denatured, His-tagged Mic40 recombinant protein as produced for LUV deformation assays.

### Mic34 site directed mutagenesis and deletion of LBS motifs

A PCR reaction was performed on the pT7-3V5-Mic34 plasmid using oligos LUK0227 and LUK0280, the former incorporating the mismatches necessary for introducing the desired point mutations within the mitofilin domain of Mic34. ΔLBS1 and ΔLBS1+ 2 deletion mutants were generated using either pT7-3V5-Mic34 or pMAL-p2X-Mic34 templates for a PCR reaction with a primer downstream of LBS1 (ΔLBS1-R) and upstream of either LBS1 (ΔLBS1-F) or LBS2 (ΔLBS2-F), respectively. Primer sequences are:

LUK0280 5’-TATCGTTGAGGCGGGGAAAGCTCCCG-3’

LUK0227 5’-GCGCGTCCGAGATTATCGGCCGTTACATCCACAACAGCTGGTGA-3’

ΔLBS1-R 5’-TGAAGGGATGCTGTTTATGTACTTGAG-3’

ΔLBS1-F 5’-GCTCCCGCGGCTGAGCCCGTTC-3’

ΔLBS2-F 5’-TTCCGCACATCACTAAGCCCAACC-3’

These oligos containing 5’phosphates were ligated using T4 ligase (NEB) following purification of the PCR product and DpnI digestion (to remove the original plasmid). The resulting plasmid was verified by sequencing.

### Immuno-electron tomography and transmission electron microscopy

The cell suspension was fixed in a mixture of 4% paraformaldehyde and 0.1% glutaraldehyde in 0.1 M HEPES buffer for 1 h at RT. The cells were washed three times in 0.1 M HEPES buffer and embedded in 10% gelatine preheated at 37°C. The gelatine blocks were immersed in 2.3 M sucrose for 72 h. The sample was placed on a stub and frozen in liquid nitrogen, then cryo-sectioned into 80-100 nm sections, using a cryo-ultramicrotome EM UC6 (Leica). The sections were placed on a formvar/carbon-coated nickel grid (300 Mesh) and allowed to thaw. The grids were then placed in 0.1 M HEPES buffer. After treatment with blocking buffer (BB) composed of 1% fish skin gelatin dissolved in HEPES for 1 h, the grids were incubated with anti-MBP antibody diluted 1:20 in BB at RT for 30 min. After washing in BB, the sections were incubated with the Protein A conjugated with 5 nm gold nanoparticles (UMC, Utrecht) diluted 1:50 in BB and incubated for 45 min at RT. Controls for nonspecific binding of the secondary antibody were performed by omitting the primary antibody. Finally, the sections were washed in HEPES and dH_2_O, stained with uranyl acetate, embedded in methylcellulose, and inspected using a JEM-1400Flash transmission electron microscope (JEOL).

Localization and analysis of immunogold labelling were performed utilizing electron tomography. A JEM 2100F (JEOL) transmission electron microscope equipped with a direct electron detector K2 Summit (Gatan) and controlled with the software package SerialEM was used to acquire tilt series images in the range +57° to -63° with a tilt step of 1°. Electron tomography reconstruction and filtering using nonlinear anisotropic diffusion were performed in the software package IMOD (Mastronarde and Held 2017).

LUVs were imaged using a Talos L120C electron microscope equipped with a Ceta 16M camera.

### Vesicle and nanoparticle counting

Particle detection was performed by applying a density threshold to enhance the contrast between the nanoparticles and the surrounding membrane. Particle counting was conducted using the ImageJ software, where the occurrence of nanoparticles and vesicles was analyzed with a precision limit of 5 nm. Vesicle counting was performed blind in 23 cells.

### Light microscopy and image analysis

Images for 2D branching analysis were acquired on a BX63 widefield microscope (Olympus) equipped with a 100× UPLSAPO NA 1.40 objective lens (Olympus), with an X-Cite excitation light source (Excelitas), U-FUNA and U-FBWA fluorescence filter cubes (Olympus), and a DP74 camera (Olympus) set to the 8-bit grayscale mode. Raw 8-bit grayscale images were converted into the 32-bit format. Images were filtered using the difference of Gaussians s = 2 and 4. Background subtraction was done in Fiji (Schindelin et al. 2012) using the sliding paraboloid and the radius of 20.0 pixels. Image thresholding was done using the triangle method. Branching quantification and analysis was performed in Fiji using the 2D analysis option of the Mitochondria Analyzer plugin (Chaudhry et al. 2019). We measured mitochondria branching and length for 100 complete cells for each condition. All statistical analyses were performed using the GraphPad Prism software, using an unpaired two-tailed Student’s t-test. A *p*-value of less than 0.05 was considered statistically significant.

Images for fragmentation analysis and 3D reconstruction were acquired on a FV3000 laser scanning confocal microscope (Olympus). The microscope was equipped with a UAPON 100× TIRF NA 1.49 objective lens (Olympus), 405 nm and 488 nm lasers for excitation (Coherent), quadband dichroic beamsplitter TRF89902 ET (Chroma), and high-sensitivity GaAsP spectral detectors with detection windows set to 420-470 nm and 500-540 nm, respectively. Z-stacks were acquired using Nyquist sampling and processed using the 3D constrained iterative deconvolution with the maximum likelihood algorithm and the theoretical point spread function in cellSens Dimension (Olympus). Creation of 3D models was done in Imaris 10.0 (Oxford Instruments) using manual thresholding for surface creation and removal of small objects. For fragmentation analysis maximum intensity projections were taken from Z-stacks and processed in Fiji using the Mitochondria Analyzer plugin. Single cells were cropped from the images and subjected to image thresholding using the default parameters in Mitochondria Analyzer with the Block size of 1.85 micron and the C-value of 3. The number of non-connected fragments for individual cells was counted from binary images. We measured 35-50 cells for each condition. All statistical analyses were performed using the GraphPad Prism software, using an unpaired two-tailed Mann-Whitney test in GraphPad Prism. A *p*-value of less than 0.05 was considered statistically significant.

## ACKOWLEDGEMENTS

We would like to thank André Schneider (University of Bern) and Alena Zíková (Biology Centre, Czech Academy of Sciences) for antibodies (AS: Tim9, Mic34; AZ: Rieske, COXIV, ATPβ, HSP70) plus Lawrence Rudy Cadena (Heinrich Heine University) for his input and Julius Lukeš (BC CAS) for his support and Erv1 antibody. This work was supported by Czech Science Foundation grant 23-07674S to HH. ME was supported by the European Union under the LERCO project number CZ.10.03.01/00/22_003/0000003 via the Operational Programme Just Transition. TB and MT were supported by the Czech BioImaging grant from the ERD Funds of the Czech Ministry of Education. BTK was supported by the Boehringer Ingelheim Fonds PhD Fellowship. BTK and MM were supported by the Deutsche Forschungsgemeinschaft (SFB-1638/1 – 511488495 – P08 and FOR-2848 – 401510699 - P05). The authors gratefully acknowledge Marek Vrbacký at the Proteomics Service Laboratory at the Institute of Physiology (supported by RVO, ID 67985823) and Institute of Molecular Genetics (supported by RVO, ID 68378050) of the Czech Academy of Sciences. The authors thank the Laboratory of Microscopy and Histology at the Biology Centre, Czech Academy of Sciences for provision of instruments. We would like to acknowledge access to the infrastructure and support provided by the Cryo-EM Network at the Heidelberg University (HDcryoNet), which is funded and supported by the German Research Founda1on (DFG), the Federal Ministry of Educa1on and Research (BMBF) and the Ministry of Science Baden-WulJrOemberg, among others, within the framework of the Excellence Strategy of the Federal and State Governments of Germany.

## DATA AVAILABILITY

The mass spectrometry proteomics data have been deposited to the ProteomeXchange Consortium via the PRIDE partner repository with the dataset identifier PXD059459. Multiple sequence alignments used in the analyses reported in the study are available in the Figshare data repository (10.6084/m9.figshare.30001090).

## REFERENCES

Aaltonen MJ, Friedman JR, Osman C, Salin B, di Rago JP, Nunnari J, Langer T, Tatsuta T. 2016. MICOS and phospholipid transfer by Ups2-Mdm35 organize membrane lipid synthesis in mitochondria. J. Cell Biol. 213:525–534.

Altschul SF, Madden TL, Schäffer AA, Zhang J, Zhang Z, Miller W, Lipman DJ. 1997. Gapped BLAST and PSI-BLAST: a new generation of protein database search programs. Nucl. Acids Res. 25:3389–3402.

Alvarez-Jarreta J, Amos B, Aurrecoechea C, Bah S, Barba M, Barreto A, Basenko EY, Belnap R, Blevins A, Böhme U, et al. 2024. VEuPathDB: the eukaryotic pathogen, vector and host bioinformatics resource center in 2023. Nucl. Acids Res. 52:D808–D816.

Barbot M, Jans DC, Schulz C, Denkert N, Kroppen B, Hoppert M, Jakobs S, Meinecke M. 2015. Mic10 oligomerizes to bend mitochondrial inner membranes at cristae junctions. Cell Metab. 21:756–763.

Barbot M, Meinecke M. 2016. Reconstitutions of mitochondrial inner membrane remodeling. J. Struct. Biol. 196:20–28.

Benning FMC, Bell TA, Nguyen TH, Syau D, Connell LB, Liao Y-T, Keating MP, Coughlin M, Nordstrom AEH, Ericsson M, et al. 2025. Ancestral sequence reconstruction of Mic60 reveals a residue signature supporting respiration in yeast. Protein Sci. 34:e70207.

Benz C, Müller N, Kaltenbrunner S, Váchová H, Vancová M, Lukeš J, Varga V, Hashimi H. 2022. Kinetoplastid-specific X2-family kinesins interact with a kinesin-like pleckstrin homology domain protein that localizes to the trypanosomal microtubule quartet. Mol. Microbiol. 118:155–174.

Bílý T, Sheikh S, Mallet A, Bastin P, Pérez-Morga D, Lukeš J, Hashimi H. 2021. Ultrastructural Changes of the Mitochondrion During the Life Cycle of *Trypanosoma brucei*. J. Eukaryot. Microbiol. 68:e12846.

Bock-Bierbaum T, Funck K, Wollweber F, Lisicki E, von der Malsburg K, von der Malsburg A, Laborenz J, Noel JK, Hessenberger M, Jungbluth S, et al. 2022. Structural insights into crista junction formation by the Mic60-Mic19 complex. Sci. Adv. 8:eabo4946.

Burki F, Roger AJ, Brown MW, Simpson AGB. 2020. The New Tree of Eukaryotes. Trends Ecol. Evol. 35:43–55.

Cadena LR, Gahura O, Panicucci B, Zíková A, Hashimi H, Blader Ira J. 2021. Mitochondrial Contact Site and Cristae Organization System and F1FO-ATP Synthase Crosstalk Is a Fundamental Property of Mitochondrial Cristae. mSphere 6:e00327–00321.

Capella-Gutiérrez S, Silla-Martínez JM, Gabaldón T. 2009. trimAl: a tool for automated alignment trimming in large-scale phylogenetic analyses. Bioinformatics 25:1972–1973.

Cavalier-Smith T. 2016. Higher classification and phylogeny of Euglenozoa. Eur. J. Protistol. 56:250–276.

Chaudhry A, Shi R, Luciani DS. 2019. A pipeline for multidimensional confocal analysis of mitochondrial morphology, function, and dynamics in pancreatic β-cells. Am. J. Physiol. Endocrinol. Metab. 318:E87–E101.

Colina-Tenorio L, Horten P, Pfanner N, Rampelt H. 2020. Shaping the mitochondrial inner membrane in health and disease. J. Intern. Med. 287:645–664.

Daumke O, van der Laan M. 2025. Molecular machineries shaping the mitochondrial inner membrane. Nat. Rev. Mol. Cell Biol.:10.1038/s41580-41025-00854-z.

Eddy SR. 2011. Accelerated Profile HMM Searches. PLoS Comput. Biol. 7:e1002195.

Eichenberger C, Oeljeklaus S, Bruggisser J, Mani J, Haenni B, Kaurov I, Niemann M, Zuber B, Lukeš J, Hashimi H, et al. 2019. The highly diverged trypanosomal MICOS complex is organized in a nonessential integral membrane and an essential peripheral module. Mol. Microbiol. 12:1731–1743.

Flaspohler John A, Jensen Bryan C, Saveria T, Kifer Charles T, Parsons M. 2010. A Novel Protein Kinase Localized to Lipid Droplets Is Required for Droplet Biogenesis in Trypanosomes. Eukaryot. Cell 9:1702–1710.

Gleisner M, Kroppen B, Fricke C, Teske N, Kliesch T-T, Janshoff A, Meinecke M, Steinem C. 2016. Epsin N-terminal Homology Domain (ENTH) Activity as a Function of Membrane Tension. J. Biol. Chem. 291:19953–19961.

Haindrich AC, Boudová M, Vancová M, Peña-Diaz P, Horáková E, Lukeš J. 2017. The intermembrane space protein Erv1 of *Trypanosoma brucei* is essential for mitochondrial Fe-S cluster assembly and operates alone. Mol. Biochem. Parasitol. 214:47–51.

Hammond MJ, Nenarokova A, Butenko A, Zoltner M, Dobáková EL, Field MC, Lukeš J. 2020. A Uniquely Complex Mitochondrial Proteome from Euglena gracilis. Mol Biol Evol 37:2173–2191.

Hashimi H. 2019. A parasite’s take on the evolutionary cell biology of MICOS. PLoS Pathog. 15:e1008166.

Hessenberger M, Zerbes RM, Rampelt H, Kunz S, Xavier AH, Purfurst B, Lilie H, Pfanner N, van der Laan M, Daumke O. 2017. Regulated membrane remodeling by Mic60 controls formation of mitochondrial crista junctions. Nat. Commun. 8:15258.

Hirumi H, Hirumi K. 1989. Continuous cultivation of Trypanosoma brucei blood stream forms in a medium containing a low concentration of serum protein without feeder cell layers. J. Parasitol. 75:985–989.

Horvath SE, Daum G. 2013. Lipids of mitochondria. Prog. Lipid Res. 52:590–614.

Huerta-Cepas J, Serra F, Bork P. 2016. ETE 3: Reconstruction, Analysis, and Visualization of Phylogenomic Data. Mol. Biol. Evol. 33:1635–1638.

Hughes L, Borrett S, Towers K, Starborg T, Vaughan S. 2017. Patterns of organelle ontogeny through a cell cycle revealed by whole-cell reconstructions using 3D electron microscopy. J. Cell Sci. 130:637–647.

Huynen MA, Muhlmeister M, Gotthardt K, Guerrero-Castillo S, Brandt U. 2016. Evolution and structural organization of the mitochondrial contact site (MICOS) complex and the mitochondrial intermembrane space bridging (MIB) complex. Biochim. Biophys. Acta 1863:91–101.

Jakob M, Hoffmann A, Amodeo S, Peitsch C, Zuber B, Ochsenreiter T. 2016. Mitochondrial growth during the cell cycle of *Trypanosoma brucei* bloodstream forms. Sci. Rep. 6:36265.

Johnston HE, Yadav K, Kirkpatrick JM, Biggs GS, Oxley D, Kramer HB, Samant RS. 2022. Solvent Precipitation SP3 (SP4) Enhances Recovery for Proteomics Sample Preparation without Magnetic Beads. Anal. Chem. 94:10320–10328.

Katoh K, Rozewicki J, Yamada KD. 2019. MAFFT online service: multiple sequence alignment, interactive sequence choice and visualization. Brief. Bioinform. 20:1160–1166.

Kaurov I, Heller J, Deisenhammer S, Potěšil D, Zdráhal Z, Hashimi H. 2022. The essential cysteines in the CIPC motif of the thioredoxin-like Trypanosoma brucei MICOS subunit TbMic20 do not form an intramolecular disulfide bridge in vivo. Mol. Biochem. Parasitol. 248:111463.

Kaurov I, Vancová M, Schimanski B, Cadena LR, Heller J, Bílý T, Potěšil D, Eichenberger C, Bruce H, Oeljeklaus S, et al. 2018. The diverged trypanosome MICOS complex as a hub for mitochondrial cristae shaping and protein import. Curr. Biol. 28:3393–3407.

Kostygov AY, Karnkowska A, Votýpka J, Tashyreva D, Maciszewski K, Yurchenko V, Lukeš J. 2021. Euglenozoa: taxonomy, diversity and ecology, symbioses and viruses. Open Biol. 11:200407.

Kroppen B, Teske N, Yambire KF, Denkert N, Mukherjee I, Tarasenko D, Jaipuria G, Zweckstetter M, Milosevic I, Steinem C, et al. 2021. Cooperativity of membrane-protein and protein–protein interactions control membrane remodeling by epsin 1 and affects clathrin-mediated endocytosis. Cell. Mol. Life Sci. 78:2355–2370.

Kühlbrandt W. 2019. Structure and mechanisms of F-type ATP synthases. Annu. Rev. Biochem. 88:515–549.

Lax G, Kolisko M, Eglit Y, Lee WJ, Yubuki N, Karnkowska A, Leander BS, Burger G, Keeling PJ, Simpson AGB. 2021. Multigene phylogenetics of euglenids based on single-cell transcriptomics of diverse phagotrophs. Mol. Phylogen. Evol. 159:107088.

Letunic I, Bork P. 2024. Interactive Tree of Life (iTOL) v6: recent updates to the phylogenetic tree display and annotation tool. Nucl. Acids Res. 52:W78–W82.

Mastronarde DN, Held SR. 2017. Automated tilt series alignment and tomographic reconstruction in IMOD. J. Struct. Biol. 197:102–113.

Meng EC, Goddard TD, Pettersen EF, Couch GS, Pearson ZJ, Morris JH, Ferrin TE. 2023. UCSF ChimeraX: Tools for structure building and analysis. Protein Sci. 32:e4792.

Michaud M, Gros V, Tardif M, Brugiere S, Ferro M, Prinz WA, Toulmay A, Mathur J, Wozny M, Falconet D, et al. 2016. AtMic60 is involved in plant mitochondria lipid trafficking and is part of a large complex. Curr. Biol. 26:627–639.

Miroux B, Walker JE. 1996. Over-production of Proteins in *Escherichia coli*: Mutant Hosts that Allow Synthesis of some Membrane Proteins and Globular Proteins at High Levels. J. Mol. Biol. 260:289–298.

Mordas A, Tokatlidis K. 2015. The MIA pathway: a key regulator of mitochondrial oxidative protein folding and biogenesis. Acc. Chem. Res. 48:2191–2199.

Muñoz-Gómez SA, Cadena LR, Gardiner AT, Leger MM, Sheikh S, Connell LB, Bilý T, Kopejtka K, Beatty JT, Koblížek M, et al. 2023. Intracytoplasmic-membrane development in alphaproteobacteria involves the homolog of the mitochondrial crista-developing protein Mic60. Curr. Biol. 33:1099–1111.e1096.

Muñoz-Gómez SA, Slamovits CH, Dacks JB, Baier KA, Spencer KD, Wideman JG. 2015. Ancient homology of the mitochondrial contact site and cristae organizing system points to an endosymbiotic origin of mitochondrial cristae. Curr. Biol. 25:1489–1495.

Muñoz-Gomez SA, Wideman JG, Roger AJ, Slamovits CH. 2017. The origin of mitochondrial cristae from alphaproteobacteria. Mol. Biol. Evol. 34:943–956.

Niemann M, Wiese S, Mani J, Chanfon A, Jackson C, Meisinger C, Warscheid B, Schneider A. 2013. Mitochondrial outer membrane proteome of Trypanosoma brucei reveals novel factors required to maintain mitochondrial morphology. Mol. Cell. Proteom. 12:515–528.

Pánek T, Eliáš M, Vancová M, Lukeš J, Hashimi H. 2020. Returning to the Fold for Lessons in Mitochondrial Crista Diversity and Evolution. Curr. Biol. 30:R575–R588.

Peikert CD, Mani J, Morgenstern M, Kaser S, Knapp B, Wenger C, Harsman A, Oeljeklaus S, Schneider A, Warscheid B. 2017. Charting organellar importomes by quantitative mass spectrometry. Nat. Commun. 8:15272.

Poon SK, Peacock L, Gibson W, Gull K, Kelly S. 2012. A modular and optimized single marker system for generating *Trypanosoma brucei* cell lines expressing T7 RNA polymerase and the tetracycline repressor. Open Biol. 2:110037.

Prokopchuk G, Butenko A, Dacks JB, Speijer D, Field MC, Lukeš J. 2023. Lessons from the deep: mechanisms behind diversification of eukaryotic protein complexes. Biol. Rev. Camb. Philos. Soc. 98:1910–1927.

Rabl R, Soubannier V, Scholz R, Vogel F, Mendl N, Vasiljev-Neumeyer A, Korner C, Jagasia R, Keil T, Baumeister W, et al. 2009. Formation of cristae and crista junctions in mitochondria depends on antagonism between Fcj1 and Su e/g. J. Cell Biol. 185:1047–1063.

Rampelt H, Wollweber F, Licheva M, de Boer R, Perschil I, Steidle L, Becker T, Bohnert M, van der Klei I, Kraft C, et al. 2022. Dual role of Mic10 in mitochondrial cristae organization and ATP synthase-linked metabolic adaptation and respiratory growth. Cell Rep. 38:110290.

Richter DJ, Berney C, Strassert JFH, Poh Y-P, Herman EK, Muñoz-Gómez SA, Wideman JG, Burki F, de Vargas C. 2022. EukProt: A database of genome-scale predicted proteins across the diversity of eukaryotes. Peer Community J. 2:e56.

Roger AJ, Munoz-Gomez SA, Kamikawa R. 2017. The origin and diversification of mitochondria. Curr. Biol. 27:1177–1192.

Schindelin J, Arganda-Carreras I, Frise E, Kaynig V, Longair M, Pietzsch T, Preibisch S, Rueden C, Saalfeld S, Schmid B, et al. 2012. Fiji: an open-source platform for biological-image analysis. Nat. Methods 9:676–682.

Smith JT, Singha UK, Misra S, Chaudhuri M. 2018. Divergent Small Tim Homologues Are Associated with TbTim17 and Critical for the Biogenesis of TbTim17 Protein Complexes in *Trypanosoma brucei*. mSphere 3:e00204–00218.

Söding J, Biegert A, Lupas AN. 2005. The HHpred interactive server for protein homology detection and structure prediction. Nucl. Acids Res. 33:W244–W248.

Stamatakis A. 2014. RAxML version 8: a tool for phylogenetic analysis and post-analysis of large phylogenies. Bioinformatics 30:1312–1313.

Stephan T, Brüser C, Deckers M, Steyer AM, Balzarotti F, Barbot M, Behr TS, Heim G, Hübner W, Ilgen P, et al. 2020. MICOS assembly controls mitochondrial inner membrane remodeling and crista junction redistribution to mediate cristae formation. EMBO J. 39:e104105.

Tarasenko D, Barbot M, Jans DC, Kroppen B, Sadowski B, Heim G, Mobius W, Jakobs S, Meinecke M. 2017. The MICOS component Mic60 displays a conserved membrane-bending activity that is necessary for normal cristae morphology. J. Cell Biol. 216:889–899.

Turra GL, Liedgens L, Sommer F, Schneider L, Zimmer D, Vilurbina Perez J, Koncarevic S, Schroda M, Muhlhaus T, Deponte M. 2021. In Vivo Structure-Function Analysis and Redox Interactomes of Leishmania tarentolae Erv. Microbiol. Spectr. 9:e0080921.

Van Laar VS, Berman SB, Hastings TG. 2016. Mic60/mitofilin overexpression alters mitochondrial dynamics and attenuates vulnerability of dopaminergic cells to dopamine and rotenone. Neurobiol. Dis. 91:247–261.

Varabyova A, Topf U, Kwiatkowska P, Wrobel L, Kaus-Drobek M, Chacinska A. 2013. Mia40 and MINOS act in parallel with Ccs1 in the biogenesis of mitochondrial Sod1. FEBS J. 280:4943–4959.

von der Malsburg K, Muller JM, Bohnert M, Oeljeklaus S, Kwiatkowska P, Becker T, Loniewska-Lwowska A, Wiese S, Rao S, Milenkovic D, et al. 2011. Dual role of mitofilin in mitochondrial membrane organization and protein biogenesis. Dev. Cell 21:694–707.

Wenger C, Oeljeklaus S, Warscheid B, Schneider A, Harsman A. 2017. A trypanosomal orthologue of an intermembrane space chaperone has a non-canonical function in biogenesis of the single mitochondrial inner membrane protein translocase. PLoS Pathog. 13:e1006550.

Wheeler RJ. 2021. A resource for improved predictions of Trypanosoma and Leishmania protein three-dimensional structure. PLoS One 16:e0259871.

Zíková A. 2022. Mitochondrial adaptations throughout the Trypanosoma brucei life cycle. J. Eukaryot. Microbiol. 69:e12911.

