## Supplementary figures for "The core MICOS complex subunit Mic60 has been substituted by two cryptic mitofilin-containing proteins in Euglenozoa"

```
Probability: 99.16%,   E-value: 2.3e-7,   Score: 98.07,   Aligned cols: 160,   Identities: 4%,   Similarity: -0.001,   Template Neff: 7.9
Q ss_pred           HHHHHHHHHHhHhhHHHHHHHHHcC---------CChHHHHHHHhc-CccCCchHHHhHhhhhc---CCCCCHHHHHHHH
Q Tb927.4.630 107 FGYTSKKLISVGLNYEALKHKMKAG---------LPFNEELSAVLK-DAESEDVNLVAAPLKYI---NSIPSPAVVRVTA   173 (301)
Q Consensus   107 ygy~SK~l~s~~L~~eaL~~kv~~G---------~PF~~EL~A~~~-Da~~~d~~~v~~~l~~~---~GvPt~~~lr~~~   173 (301)
                    ....+.+.+.+|+++.+|+.++.+| .||..+|.+++. .++|+.+..+++.|++. .||||..+|+.+|
T Consensus   386 ~~~~~~~~~~l~~a~~~L~~al~~g~~~~~~~~~~~p~~~el~~l~~~~~~d~~v~~~l~~l~~~a~~~Gv~t~~~L~~rF   465 (579)
T PF09731.14  386 SELEKRAITQLTQCIVDLQKLLHDKSAIDGANGKGLLVRSFKNLTDLSNYDQLIKELLSTLPEGFEKKPVIPLDTLNNQF   465 (579)
T ss_pred           HHHHHHHHHHHHHHHHHHHHHHHhcCCcCCCccCCcchHHHHHHHHHhcCCCHHHHHHHhcCChhHhcCCCCCHHHHHHHH

Q ss_pred           HHHHHHHHHhCCC-CCCCCc--cc---hhhhhccccCCChhHHhHHHHHHHHHHHHHHHHHHHHcccHHHHHHHHHHHHH-H
Q Tb927.4.630 174 FNLGRAIVEAGKA-PAAEPV--RG---ILDFFKFRTSLSPTMLVSRQLRALKSEAEFLRFVERGEYHRALSLAEDVYA-G   246 (301)
Q Consensus   174 ~~La~ai~~~~~~-p~~~~~--r~---Wld~~kf~t~~sp~~~~~~~~~~~~~~~~~~~~l~~~e~~~Al~~~~~~~~-~   246 (301)
                    ..+++.+++++.+ |+++|+ ++ +.|+|.|++.+.+ .+.++.++|+|+.++|.+|||..|+++++.|+| +
T Consensus   466 ~~v~~~~~~~~a~~~~~~~g~~~~~~s~l~s~~~v~~~~~~-----~~~~~~~iL~ra~~~L~~gdl~~Av~~l~~L~g~~   540 (579)
T PF09731.14  466 QDIAKKLRKSQLIDPNDNSLLGKAVSELASLFIIPEKGMV-----QGNDYDAILARAEDHLRKNNLSSAIKEMESIQQQS   540 (579)
T ss_pred           HHHHHHHHHHhcCCCCCCCHHHHHHHHHHHHHhcCcccCCC-----CCCCHHHHHHHHHHHHHHCCCHHHHHHHHHcCCchh

Q ss_pred           HHhcCCcchHHHH-----------HHHHHHHHhHHHHHHHHHHHHHHH
Q Tb927.4.630 247 LKREGVKDAAVLD-----------EAFGAFRRTVLPTVAGDMLLRYC   282 (301)
Q Consensus   247 ~~~~~d~~~~~~~-----------~~~~~fr~~~~~~iaa~q~l~y~   282 (301)
                    ++ ..+.+|+.+++.|++++++++.|
T Consensus   541 -----------~~~~~~~~~~~~~~~~a~dWl~~ar~rle~~qa~~~L   576 (579)
T PF09731.14  541 -----------SKSSTNIDNHHLSKLTSNWIKQAKERDQLENISKLL   576 (579)
T ss_pred           -----------hccccCCcchhHHHHHHHHHHHHHHHHHHHHHHHHh
```

**Supplementary fig. S1. Alignment of euglenozoan Mic34 and the Pfam family PF09731.14 (Mitofilin) as built by HHpred.**

The figure shows an output of a HHpred search matching the query, i.e. a profile HMM derived from a custom alignment of euglenozoan Mic34 sequences (in the alignment represented by Mic34 from *Trypanosoma brucei*, Tb927.4.630), to a target, i.e. a profile HMM derived from PF09731.14. Significance values of the hit are presented above the alignment.

```
Probability: 98.03%,   E-value: 0.021,   Score: 63.64,   Aligned cols: 250,   Identities: 11%,   Similarity: 0.112,   Template Neff: 7.9
Q ss_pred              HHHHHHHHHHHHHHHHHHHHHHH-hhHHHHHHHHHHHHHHHHHHHHHHHHHHHHHHHHHHHHHHHHHHHHHHHHHHHHHH
Q Tb927.8.580 29       GKLRSRRADLDRQYADSLAETS-ALDEARLAGAELLRQKRKESVLLHEAMDSIWKDRFERYKQTNKETHSYLRALPEALG  107 (355)
Q Consensus   29       ~~i~~rr~~L~~E~~~l~~el~-~~~~~~~~~~el~~~~~~~~~~~~e~~~~LW~DRl~Ry~~~~~~~~~y~~alPe~l~  107 (355)
                       +.....+.++.++...++++.+ .+..+..+..+.+.+++++. ..+....+=.+| ..+...|-...+
T Consensus   304      ~~~~~~~~~~~~~~~~~~~~~~~~l~~~L~~~~~~~~~~l~~~--~~~~~~~v~~er-----------~~~~~~l~~l~~  370 (579)
T PF09731.14  304      KLINGKDDELKRVEEEIREKYRSSLDAAIEELNKDLDEKLKDM--DQFIKSKVFDNQ-----------AVLQETLEKQKS  370 (579)
T ss_pred              HHHhcHHHHHHHHHHHHHHHHHHHHHHHHHHHHHHHHHHHHHH--HHHHHHHHHHHH-----------HHHHHHHHHHHH

Q ss_pred              H-HHHHHHHHHHHHHHHHHhhcHHHHHHHHHHHHHHHcccch-------hcHHHHHHHHHHHhhcCHHHHHHHHHHHhhc
Q Tb927.8.580 108      V-LKGVTNHYQYMAREMRKFIGFDVACSKVHNFALLLNHGEK-------VGIQRVSETIRQLLVAEPLVRVVCDSLLDEA  179 (355)
Q Consensus   108      ~-l~~L~~hY~~m~~~m~~~~~fd~~~sKvH~laLll~~~~~-------~~~~~v~~~lr~lf~~DPLv~~v~~S~~~~~  179 (355)
                       . +++|.....-..........-.....=+..|.=.+..+.. ..+...+..|+.+...||+|..+..++....
T Consensus   371      ~~l~~le~~~~~~~~~~~~~~~~~~~l~~a~~~L~~al~~g~~~~~~~~~~p~~~el~~l~~~~~~d~~v~~~l~~l~~~a  450 (579)
T PF09731.14  371      NLINIFKQQAESIKQSELEKRAITQLTQCIVDLQKLLHDKSAIDGANGKGLLVRSFKNLTDLSNYDQLIKELLSTLPEGF  450 (579)
T ss_pred              HHHHHHHHHHHHHhhHHHHHHHHHHHHHHHHHHHHHHHHhcCCcCCCccCCcchHHHHHHHHHhcCCCHHHHHHHhcCChhH

Q ss_pred              cCCCCCCcHHHHHHHHHHHHHHHHHHHHHHHHHHHHhcCCCCCCCCCcHHHHHHHHHHHHhHhhccc-----hhhHHHHH
Q Tb927.8.580 180      TNFSCPTSISESSDSFVFCMEGLDRAVDAAAARYIELQKPSMGESPPNVFCRVLRKITTALEANMLS-----RCDIIMKE  254 (355)
Q Consensus   180      ~~~~~P~s~~e~s~sf~~~l~~L~~~~~~~~~~~~~~~~~~~~~~~~~~~~~~~~~~~~~~~r~~~~~-----~~~~~l~~  254 (355)
                       .....|+ ...|...-..+..........| +..+|++...++.+.+.+.+.... .....|.+
T Consensus   451      ~~~Gv~t------------~~~L~~rF~~v~~~~~~~~a~~~--~~~~g~~~~~~s~l~s~~~v~~~~~~~~~~~~~iL~r  516 (579)
T PF09731.14  451      EKKPVIP------------LDTLNNQFQDIAKKLRKSQLID--PNDNSLLGKAVSELASLFIIPEKGMVQGNDYDAILAR  516 (579)
T ss_pred              hcCCCCC------------HHHHHHHHHHHHHHHHHhhcCC--CCCCCHHHHHHHHHHHHhcCcccCCCCCCCHHHHHHH

Q ss_pred              HHHHHHHHHHHHhHHHhccHHHHHHHHHHHHHHHHHhhccCCCCcccccccCCHHHHHHH-----------HHHHHHHHH
Q Tb927.8.580 255      ERQALRNHLLRDRRQLHTMHDLVGAMRYVEELSRHFQDSAKNPASVNMSVLSDPEVLLAR-----------EQLLLWRRA  323 (355)
Q Consensus   255      ~~~~~~~~~~~~~~~l~t~~Dv~aAl~y~e~~~~~~~~~~~~~~~~~~~~~~~~~~ev~~A~-----------~qL~~W~~~  323 (355)
                       .+..+.+- |+..|+++.+.+ .+..+. .-+..|...
T Consensus   517      a~~~L~~g------------dl~~Av~~l~~L----------------------~g~~~~~~~~~~~~~~~~~~~a~dWl~~  562 (579)
T PF09731.14  517      AEDHLRKN------------NLSSAIKEMESI----------------------QQQSSKSSTNIDNHHLSKLTSNWIKQ  562 (579)
T ss_pred              HHHHHHCC------------CHHHHHHHHHcC----------------------CchhhcccccCCcchhHHHHHHHHHH

Q ss_pred              HHHHHHHHHHHHHHHH
Q Tb927.8.580 324      AAIFLVRQQAKEALNS  339 (355)
Q Consensus   324      A~~yl~~~Qa~~aL~a  339 (355)
                       |..||..+|+.++|+|
T Consensus   563      ar~rle~~qa~~~L~a  578 (579)
T PF09731.14  563      AKERDQLENISKLLEL  578 (579)
T ss_pred              HHHHHHHHHHHHHhhc
```

**Supplementary fig. S2. Alignment of euglenozoan Mic40 and the Pfam family PF09731.14 (Mitofilin) as built by HHpred.**

The figure is analogous to that shown as supplementary fig. S1, except for the query being a profile HMM derived from a custom alignment of euglenozoan Mic40 sequences (in the alignment represented by Mic34 from *Trypanosoma brucei*, Tb927.8.580).

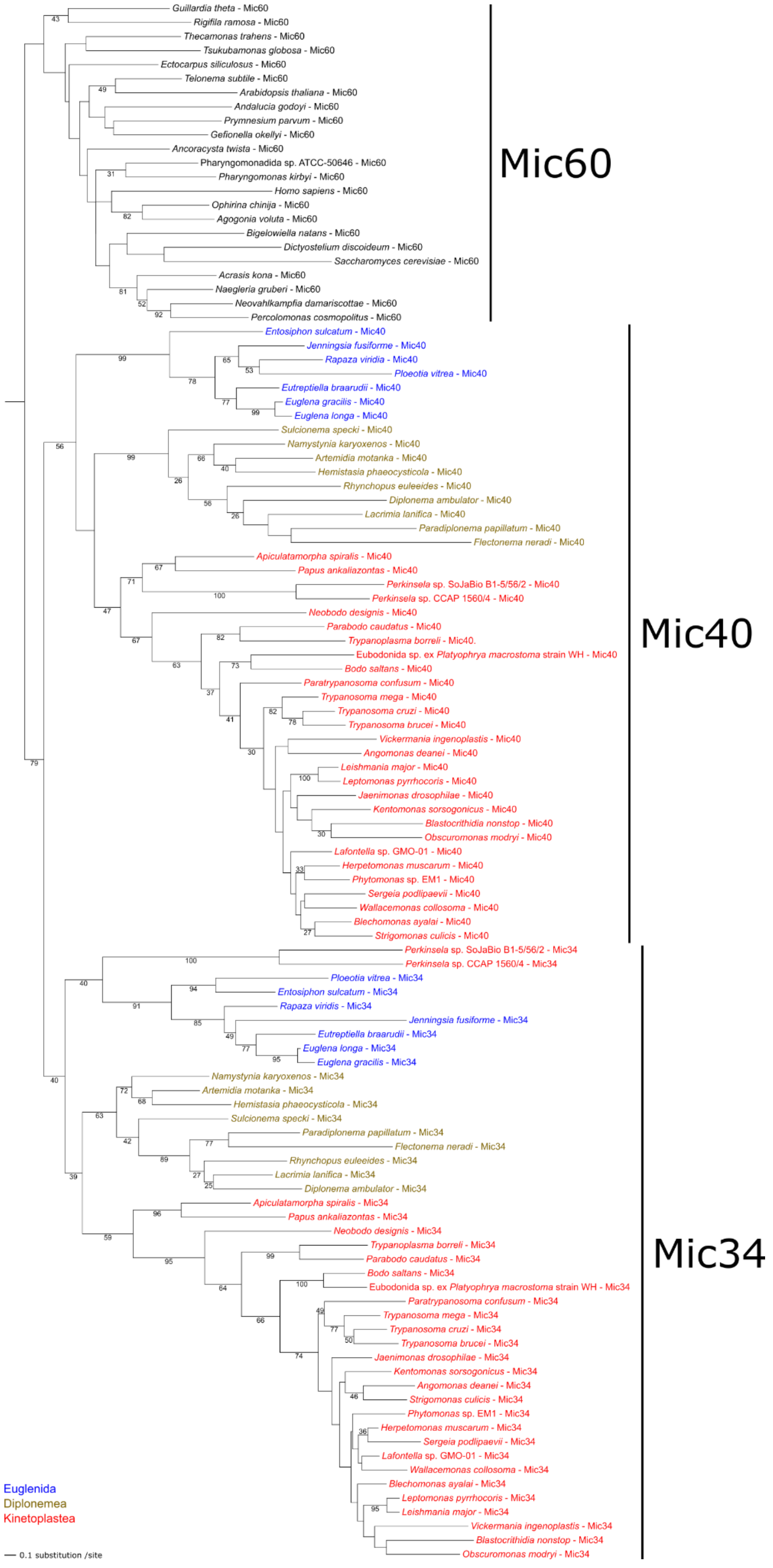

Guillardia theta - Mic60
Rigifila ramosa - Mic60
Thecamonas trahens - Mic60
Tsukubamonas globosa - Mic60
Ectocarpus siliculosus - Mic60
Telonema subtile - Mic60
Arabidopsis thaliana - Mic60
Andalucia godoyi - Mic60
Prymnesium parvum - Mic60
Gefionella okellyi - Mic60
Ancoracysta twista - Mic60
Pharyngomonadida sp. ATCC-50646 - Mic60
Pharyngomonas kirbyi - Mic60
Homo sapiens - Mic60
Ophirina chinija - Mic60
Agogonia voluta - Mic60
Bigelowiella natans - Mic60
Dictyostelium discoideum - Mic60
Saccharomyces cerevisiae - Mic60
Acrasis kona - Mic60
Naegleria gruberi - Mic60
Neovahlkampfia damariscottae - Mic60
Percolomonas cosmopolitus - Mic60
Mic60
Entosiphon sulcatum - Mic40
Jenningsia fusiforme - Mic40
Rapaza viridia - Mic40
Ploeotia vitrea - Mic40
Eutreptiella braarudii - Mic40
Euglena gracilis - Mic40
Euglena longa - Mic40
Sulcionema specki - Mic40
Namystynia karyoxenos - Mic40
Artemidia motanka - Mic40
Hemistasia phaeocysticola - Mic40
Rhynchopus euleeides - Mic40
Diplonema ambulator - Mic40
Lacrimia lanifica - Mic40
Paradiplonema papillatum - Mic40
Flectonema neradi - Mic40
Apiculatamorpha spiralis - Mic40
Papus ankaliazontas - Mic40
Perkinsela sp. SoJaBio B1-5/56/2 - Mic40
Perkinsela sp. CCAP 1560/4 - Mic40
Neobodo designis - Mic40
Parabodo caudatus - Mic40
Trypanoplasma borreli - Mic40.
Eubodonida sp. ex Platyophrya macrostoma strain WH - Mic40
Bodo saltans - Mic40
Paratrypanosoma confusum - Mic40
Trypanosoma mega - Mic40
Trypanosoma cruzi - Mic40
Trypanosoma brucei - Mic40
Vickermania ingenoplastis - Mic40
Angomonas deanei - Mic40
Leishmania major - Mic40
Leptomonas pyrrhocoris - Mic40
Jaenimonas drosophilae - Mic40
Kentomonas sorsogonicus - Mic40
Blastocrithidia nonstop - Mic40
Obscuromonas modryi - Mic40
Lafontella sp. GMO-01 - Mic40
Herpetomonas muscarum - Mic40
Phytomonas sp. EM1 - Mic40
Sergeia podlipaevii - Mic40
Wallacemonas collosoma - Mic40
Blechomonas ayalai - Mic40
Strigomonas culicis - Mic40
Mic40
Perkinsela sp. SoJaBio B1-5/56/2 - Mic34
Perkinsela sp. CCAP 1560/4 - Mic34
Ploeotia vitrea - Mic34
Entosiphon sulcatum - Mic34
Rapaza viridis - Mic34
Jenningsia fusiforme - Mic34
Eutreptiella braarudii - Mic34
Euglena longa - Mic34
Euglena gracilis - Mic34
Namystynia karyoxenos - Mic34
Artemidia motanka - Mic34
Hemistasia phaeocysticola - Mic34
Sulcionema specki - Mic34
Paradiplonema papillatum - Mic34
Flectonema neradi - Mic34
Rhynchopus euleeides - Mic34
Lacrimia lanifica - Mic34
Diplonema ambulator - Mic34
Apiculatamorpha spiralis - Mic34
Papus ankaliazontas - Mic34
Neobodo designis - Mic34
Trypanoplasma borreli - Mic34
Parabodo caudatus - Mic34
Bodo saltans - Mic34
Eubodonida sp. ex Platyophrya macrostoma strain WH - Mic34
Paratrypanosoma confusum - Mic34
Trypanosoma mega - Mic34
Trypanosoma cruzi - Mic34
Trypanosoma brucei - Mic34
Jaenimonas drosophilae - Mic34
Kentomonas sorsogonicus - Mic34
Angomonas deanei - Mic34
Strigomonas culicis - Mic34
Phytomonas sp. EM1 - Mic34
Herpetomonas muscarum - Mic34
Sergeia podlipaevii - Mic34
Lafontella sp. GMO-01 - Mic34
Wallacemonas collosoma - Mic34
Blechomonas ayalai - Mic34
Leptomonas pyrrhocoris - Mic34
Leishmania major - Mic34
Vickermania ingenoplastis - Mic34
Blastocrithidia nonstop - Mic34
Obscuromonas modryi - Mic34
Mic34
Euglenida
Diplonemea
Kinetoplastea
0.1 substitution /site

**Supplementary fig. S3. Maximum likelihood tree of the mitofilin domain.**
A tree inferred with RAxML v8.2.11 (substitution model PROTGAMMAJTT and default parameters) from a multiple alignment of 111 sequences (233 positions after trimming). Sequences representing the mitofilin domain in Mic60 proteins from various eukaryotes (with the sampling focused on representatives of the supergroup Discoba) and Mic34 and Mic40 proteins from Euglenozoa. The root was arbitrarily placed between the Mic60 and Mic34+Mic40 subtrees. Numbers at branches correspond to bootstrap values calculated from 100 replicates (shown only when ≥25%). Tip labels corresponding to euglenozoan sequences are rendered in different colors according to the classification of the source organism into one of the three main euglenozoan subgroups (see the legend on the bottom left). Sequence accession numbers are provided in supplementary dataset S1, Supplementary Material online.

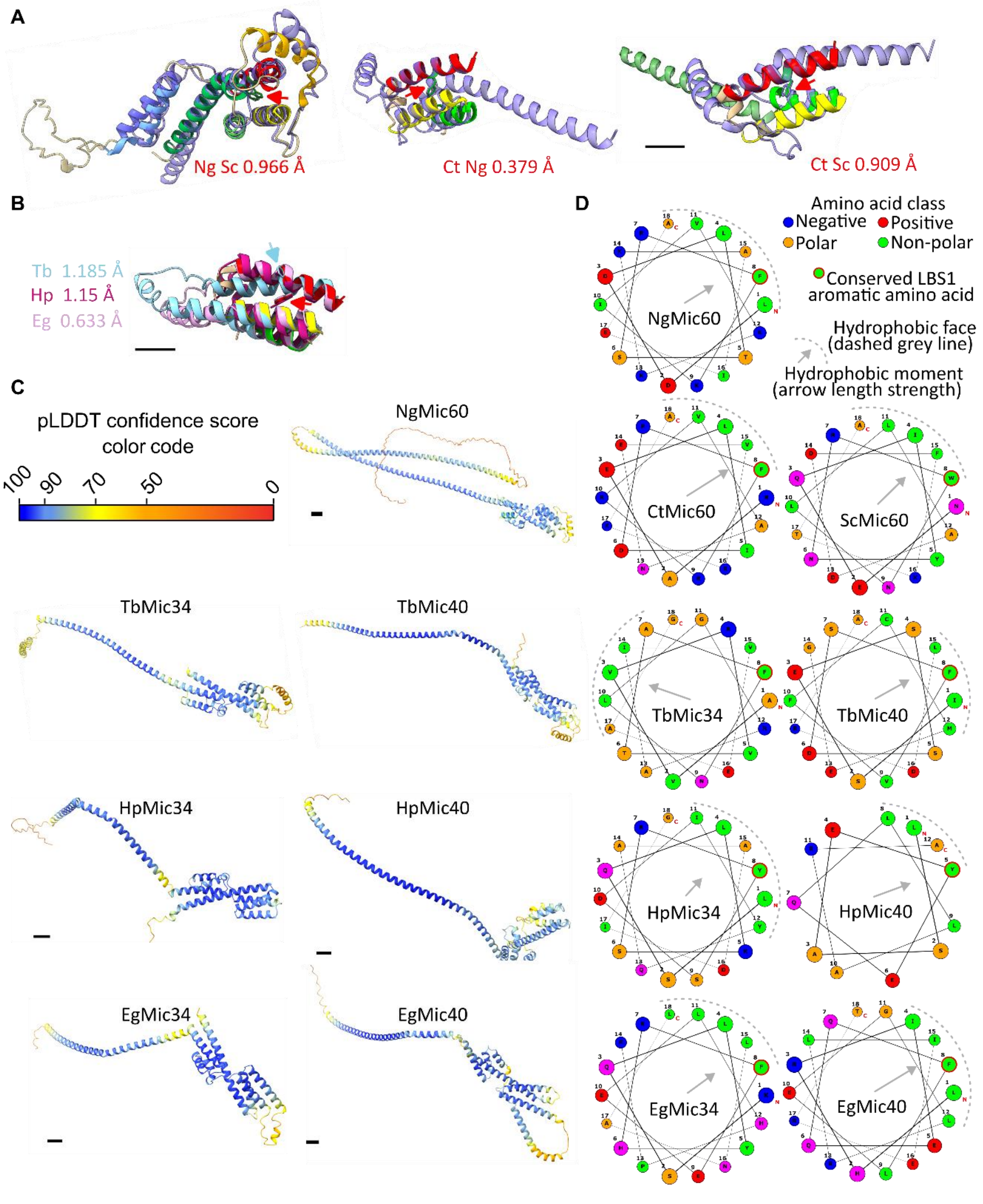


**Supplementary fig. S4. Mitofilin family structures.**

**(A)** Structural alignments of Mic60 mitofilin domains from yeast and the discoban *N. gruberi* (RDSM values given below in red). Red arrows point to aligned LBS1 aromatic amino acids. Alpha helix α8 not shown for the *Thermochaetoides thermophila* Mic60 (CtMic60). Scale bar, 10 Å.

**(B)** Structural alignment of three representative Mic34 mitofilin domains (colored as in names on the left) aligned to that of *N. gruberi* Mic60 (alpha helices colored as in supplementary fig. 1B). RMSD scores for alignment using each representative Mic34 given on left. Red arrow points to superposition of LBS1 aromatic amino acids; light blue arrow points to askew aromatic amino acid in *T. brucei* (Tb) Mic60. Scale bar, 10 Å.

**(C)** Predicted AlphaFold2 of full-length *N. gruberi* (Ng) Mic60 plus representative Mic34 and 40 depicted in supplementary fig. 1. Structures are colored according to pLDDT confidence score color coding on top left. Note that LBS2 is the least confidently predicted mitofilin alpha helix.
**(D)** Helical wheel projections of the LBS1 amphipathic alpha helices of representative Mic60, Mic34 and Mic40 structures. Key to depicted features in the projections are given in the top left.
All abbreviations as defined in the legend of Fig. 1C.

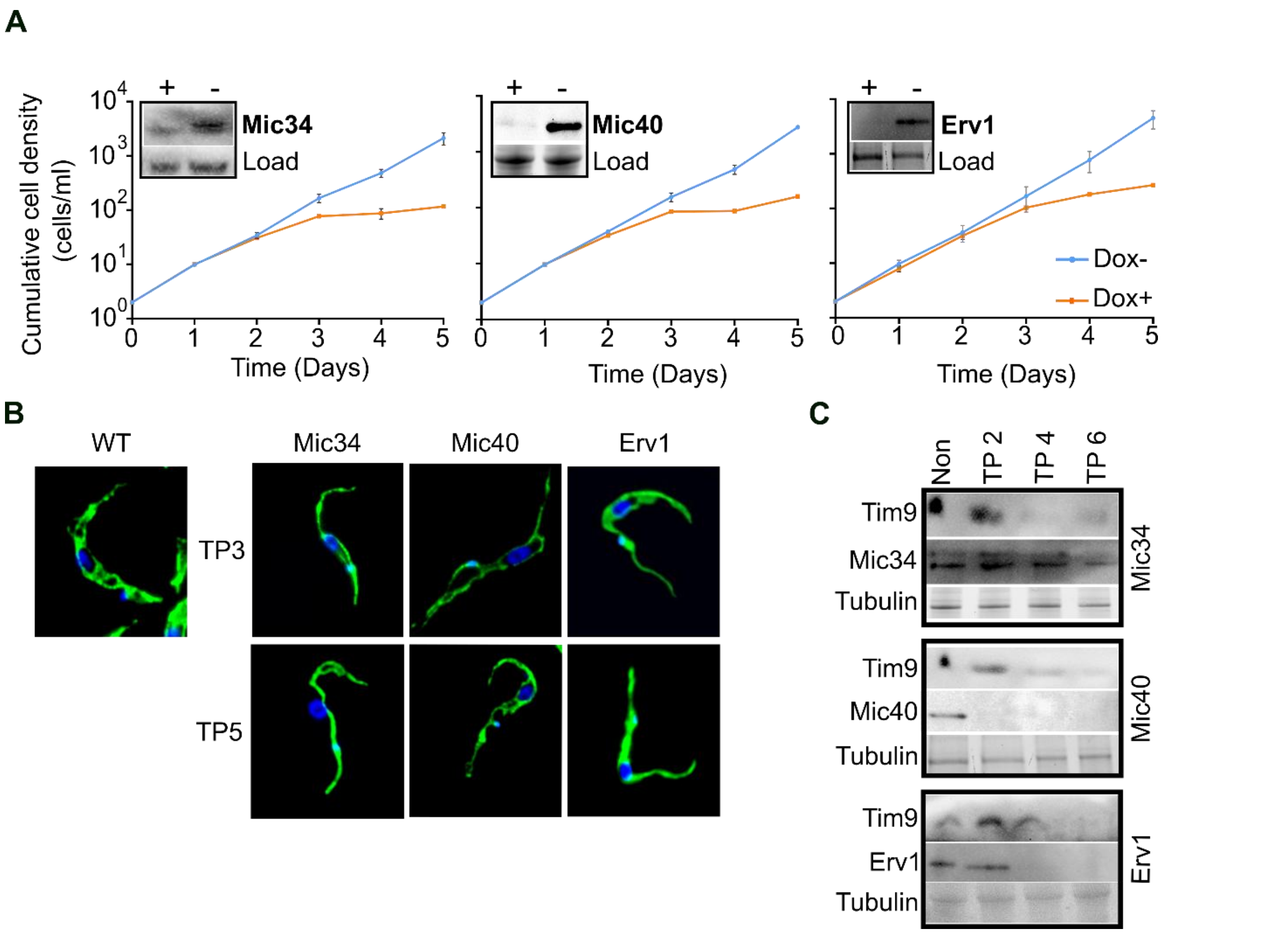


**Supplementary fig. S5. Mic34 and Mic40 depletion leads to defects in intermembrane space protein import that are associated with aberrant mitochondrial morphology, phenocopying Erv1 RNAi.**
**(A)** Accumulated cell growth measurements of Mic34, Mic40, and Erv1 RNAi for 5 days of doxycycline (Dox) induction (+) as compared to non-induced cells (-). Insets show western blots probed with an antibody recognizing Mic34, Erv1, and the HA epitope attached to Mic40 expressed from one allele. The loading control for both Mic40 and Erv1 is a fluorescently detected protein on a stain-free gel, while for Mic34 an anti-Mic34 antibody nonspecific band was used as a loading control.
**(B)** Gross morphology of mitochondria visualized by confocal microscopy of wild-type (WT) as well as Mic34, Mic40, and Erv1 knockdown cells, probed with mtHSP70 (green) and DAPI (blue), were taken at 3- and 5-days post-induction. Scale bar, 5 µm.
**(C)** Mic34, Mic40, and Erv1 RNAi cells were collected without Dox induction (Non) and at 2 (TP2), 4 (TP4), and 6 (TP6) days post-induction and proteins indicated on left were detected on western blots with specific antibodies.

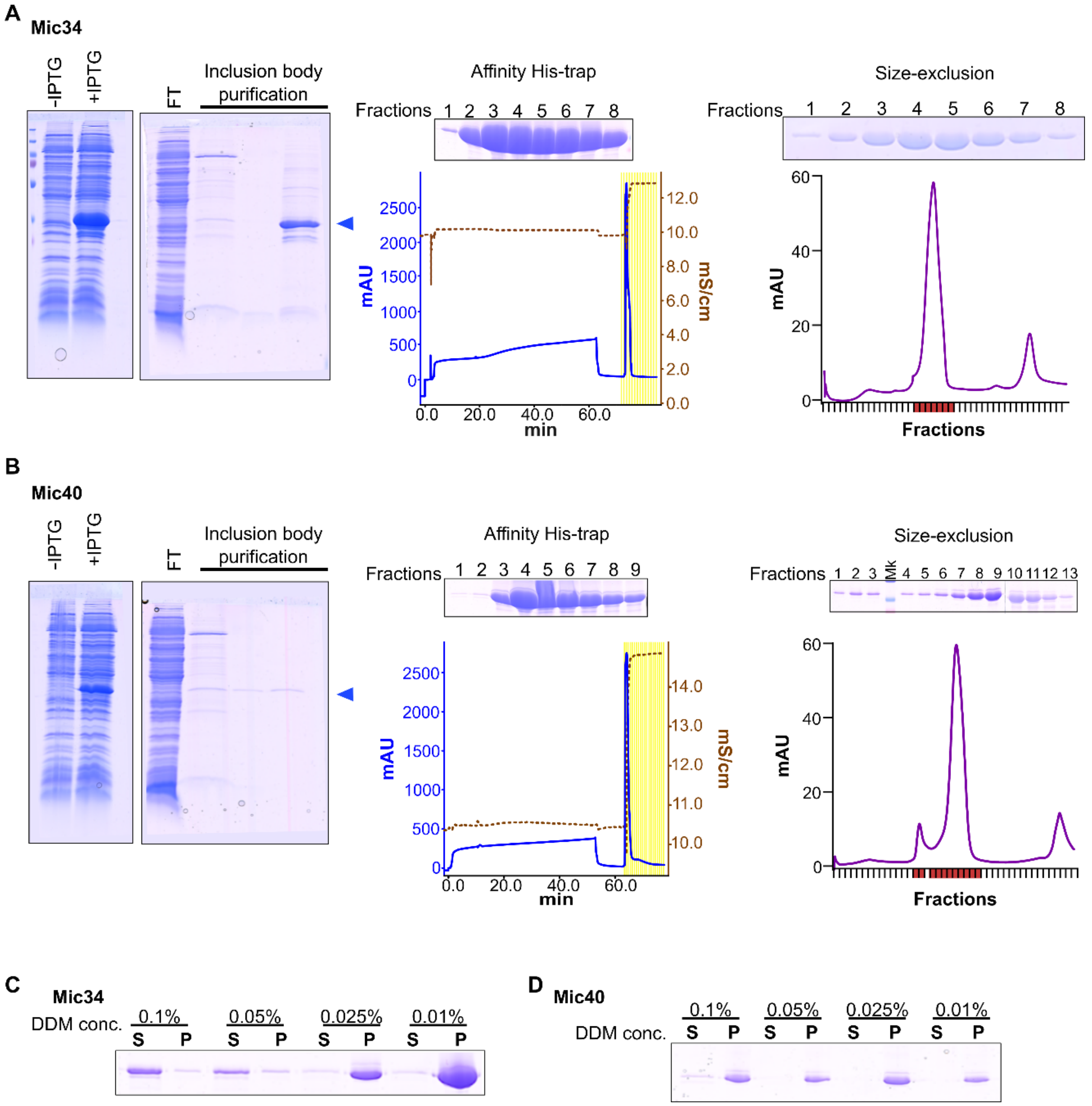


**Supplementary fig. S6. Expression and purification Mic34 and Mic40.**
**(A)** Expression and purification of recombinant Mic34 resolved by SDS-PAGE with subsequent Coomassie Brilliant Blue staining from inclusion bodies (left) by His-trap affinity chromatography and size exclusion chromatography (right). Blue arrowheads to the right of gels point to recombinant protein. The affinity His-trap chromatogram with protein absorbance measured at 280 nm in milli-absorbance units (mAU) represented by blue line and conductivity in mS/cm represented by brown dashed line. The highlighted yellow region toward the end of the chromatogram shows the elution peak. The size exclusion chromatogram shows UV absorbance at 280 nm in mAU. The peak fractions collected for SDS-PAGE analysis are highlighted in red at the base of the chromatogram.
**(B)** Expression and purification of recombinant Mic40. Data presented as in A.
**(C)** DDM concentration titration to determine appropriate Mic34 solubilization conditions as assayed by ultracentrifugation sedimentation. S, supernatant; P, pellet.
**(D)** DDM concentration titration to determine appropriate Mic40 solubilization conditions as in C.

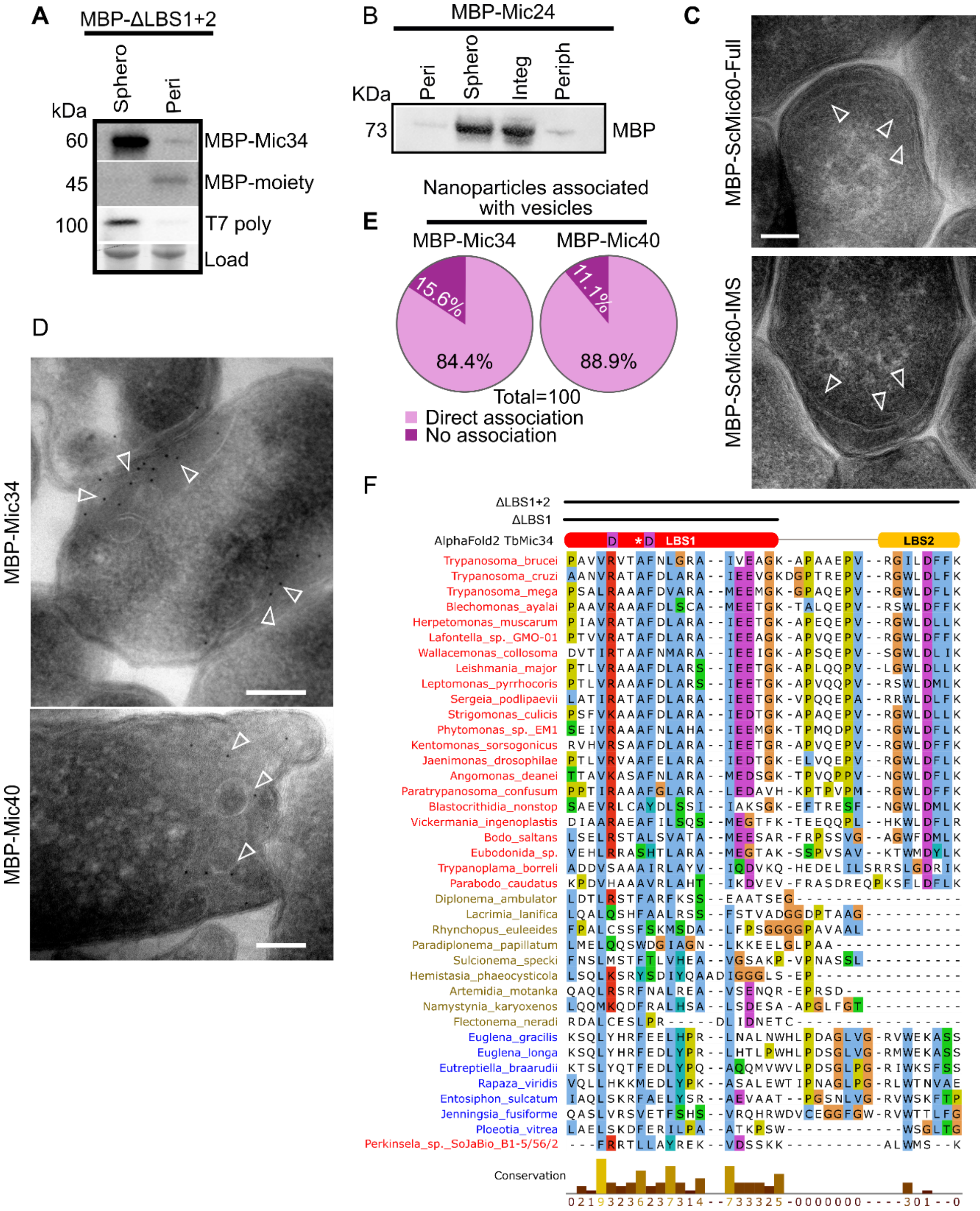


**Supplementary fig. S7. Deformation of *E. coli* cytoplasmic membranes by periplasm-targeted mitofilin family proteins.**

**(A)** Osmotic shock of MBP-Mic34ΔLBS1+2 to separate spheroplasts (Sphero) and periplasm (Peri). Western blot analysis was performed using MBP antibody and T7 polymerase (T7 Poly) as markers. A fluorescently detected protein visualized using a stain-free gel served as a loading control.

**(B)** Fractionation and carbonate extraction of MBP-Mic24 into spheroplasts (Sphero), periplasm (Peri), integral (Integ) and peripheral (Periph) proteins.

**(C)** TEM images of *E. coli* expressing full-length MBP-ScMic60-Full and soluble truncation MBP-ScMic60-IMS (9). White arrowheads point to vesicles and other deformations of the cytoplasmic membrane. Scale bar, 200 nm.
**(D)** TEM images of MBP-Mic34 and MBP-Mic40, with white arrowheads pointing to nanoparticles near vesicles. Scale bar, 200 nm.
**(E)** Pie chart quantifying nanoparticles associated with 100 vesicles in Mic34-MBP and Mic40-MBP in 4 cells observed by electron tomography.
**(F)** Multisequence alignment (MSA) of euglenozoan Mic34 LBS1 and 2. Species are given on left and colored based on euglenozoan clade: Kinetoplastea, red; Diplonemea, brown; Euglenida, blue. Schema of *T. brucei* LBS alpha helices predicted by AlphaFold2 (Fig. 1, supplementary fig. 4) is immediately above MSA. Deletions made in ΔLBS mutants (left) are indicated by uppermost black bars. Converted amino acids in TbMic34$^{R170D_F174}$ point mutants are indicated with purple-boxed 'D's, with second one marking the conserved phenylalanine (F) of kinetoplastid, while the asterisk marks conserved F position in diplonemids and euglenids.

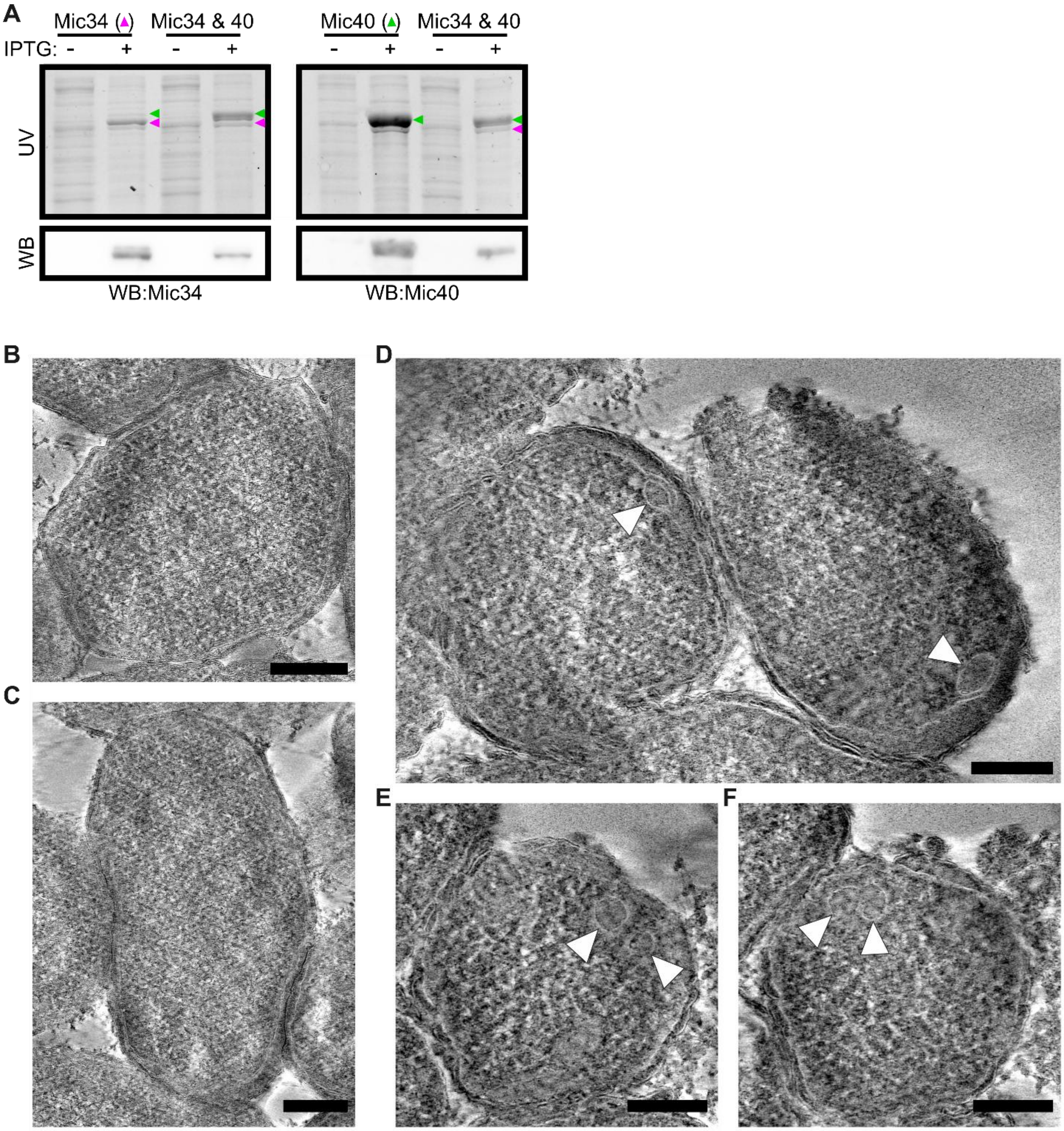


**Supplementary fig. S8. Co-expression of Mic34 and Mic40 in *E. coli* induces prominent remodeling of the cytoplasmic membrane.**
**(A)** Verification of co-expression of Mic34-MBP and Mic40-MBP in *E. coli* induced by IPTG (+) in comparison to non-induced cell (-). Top panel shows abundant proteins in the *E. coli* lysate as visualized by UV excitation of abundant proteins (see Materials and Methods). Arrowhead points to 75.1 kDa Mic34-MBP (purple) and 80.4 kDa Mic34-MBP (green). Bottom panel shows corresponding western blots of the corresponding gels in the top panel immunolabelled with antibody against Mic34 or Mic40.
(**B-C).** Electron tomograms of wild-type *E. coli*.
**(D-F)** Electron tomograms of *E. coli* co-expressing Mic34-MBP and Mic40-MBP. White arrowheads point to cytosolic vesicles, surrounded negatively stained membranes are well. The lefthand cell in D is shown in E and F at different rotation planes at increments representing ~47 nm thickness. Scale bars, 200 nm.

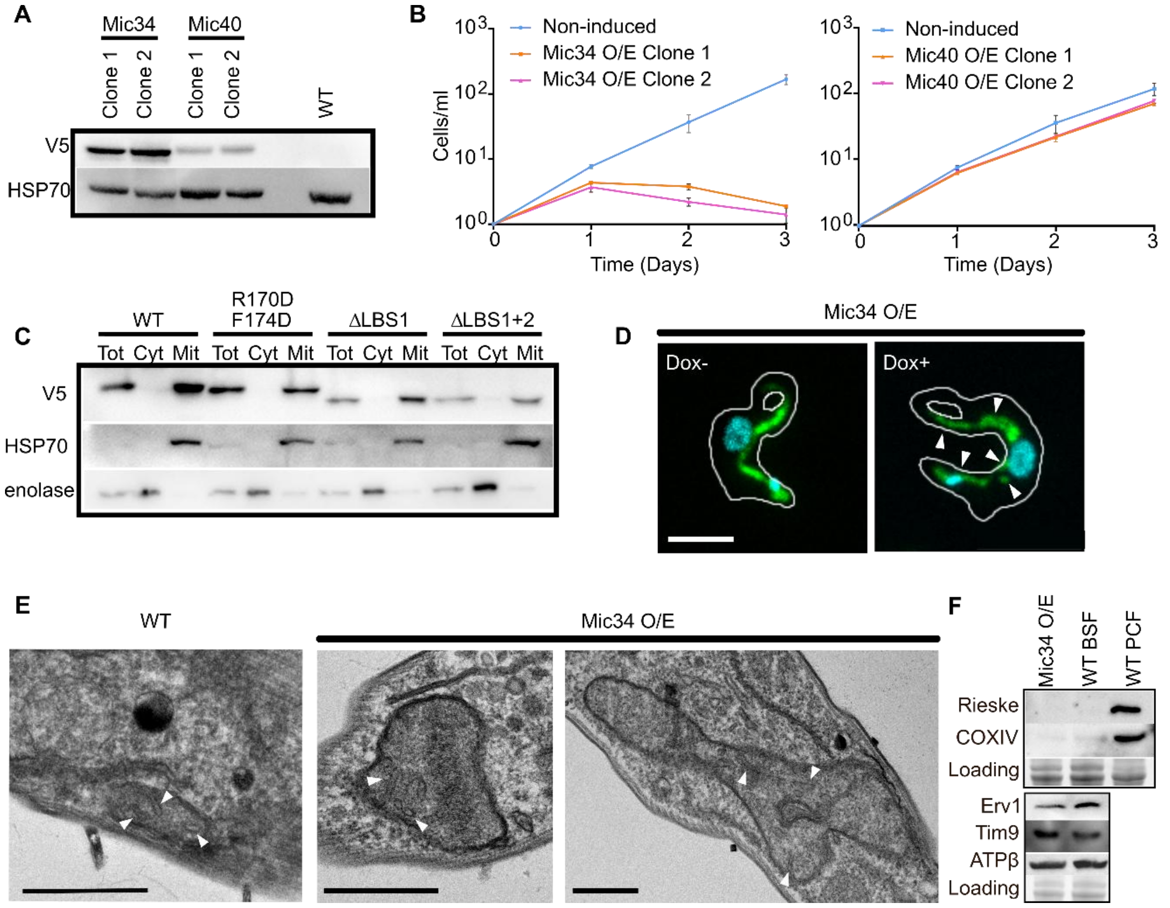


**Supplementary fig. S9. Mic34 overexpression causes growth inhibition without affecting protein import, OXPHOS complexes, or cristae morphology**.

**(A)** Western blot analysis of Mic34 and Mic40 overexpression (O/E) in two clones and WT. Blots were probed with antibodies against V5 and mitochondrial HSP70.

**(B)** Cumulative cell growth measurements of Mic34 and Mic40 O/E following three days of Dox induction. Non-induced samples are represented by a blue line, Clone 1 by an orange line, and Clone 2 by a pink line. Whiskers represent standard deviations.

**(C)** Digitonin fractionation of total *T. brucei* bloodstream form cells (Tot) overexpressing examined forms of V5-tagged Mic34 (labelled on top as in supplementary fig. 5) into cytosolic (Cyt) and mitochondrial fractions (Mit). Anti-HSP70 antibody marks fractions containing mitochondria, whereas the anti-enolase antibody marks cytosolic fractions. Note that the V5 signal fractionates with HSP70's, indicating mitochondrial targeting.

**(D)** Merged Z-stack confocal slices of *T. brucei* overexpressing Mic34 (Dox+), and non-induced control (Dox-). Cell body and flagellum outlined. Green signal is from anti-HSP70 antibody, which labels a protein dispersed throughout the mitochondrial matrix. Blue signal corresponds to stained DNA. Arrowheads point to mitochondrial fragments observed in the Mic34 overexpressing cells. Scale bar, 5 µm.

**(E)** TEM images of Mic34 O/E after 2 days of Dox induction compared to WT. White arrowheads indicate cristae structures. Scale bar, 500 nm.

**(F)** Western blot analysis of Mic34 O/E, WT BSF, and WT PCF. Antibodies used are indicated on the left of the blot.

**Supplementary movie S1 (separate file). Electron tomogram of MBP-Mic34 deformation of the *E. coli* cytoplasmic membrane.** Same cell as in Supplementary fig. 4C. Red trace, outer membrane; purple trace, cytoplasmic membrane; orange trace, vesicles; green circles, gold nanoparticles that visualize the MBP moiety of the chimeric protein**.**

**Supplementary movie S2 (separate file). Electron tomogram of MBP-Mic40 deformation of the E. coli cytoplasmic membrane.** See supplementary movie S1 legend.

**Supplementary dataset S1 (separate file).** Mic34, Mic40, Mic24, and Mic60 sequences analyzed in this study.
